# Macrophage activation of cGAS and TRIM5 distinguish pandemic and non-pandemic HIV

**DOI:** 10.1101/2022.01.21.477263

**Authors:** Lorena Zuliani Alvarez, Morten L. Govasli, Jane Rasaiyaah, Chris Monit, Stephen O. Perry, Rebecca P. Sumner, Simon McAlpine-Scott, Claire Dickson, K. M. Rifat Faysal, Laura Hilditch, Richard J. Miles, Frederic Bibollet-Ruche, Beatrice H. Hahn, Till Boecking, Nikos Pinotsis, Leo C. James, David A. Jacques, Greg J. Towers

**Author notes:** Correspondance David A. Jacques and Greg J. Towers.

## Abstract

Pandemic viruses remain a global threat to health and economics but how they adapt to become pandemic remains poorly understood. Here we compare pandemic HIV-1(M) and non-pandemic HIV-(O) and HIV-2 strains finding that non-pandemic HIV replicate poorly in myeloid cell models due to activation of cGAS and TRIM5, and ensuing antiviral responses. We use phylogenetics and viral capsid structural biology to define specific differences between pandemic and non-pandemic HIV capsids and demonstrate that their genetic reversal in HIV-1(M) mutants causes TRIM5, cGAS and innate immune activation. We propose a model in which the parental lineage of pandemic HIV-1(M) has uniquely evolved a dynamic capsid that avoids activation of cGAS and TRIM5 to establish cloaked replication in myeloid cells. The unique adaptations of the pandemic virus lineage suggests a role in effective human-to-human transmissibility and highlight the importance of avoiding innate immune activation during pandemic human-to-human viral transmission.

## INTRODUCTION

Human immunodeficiency viruses (HIVs) have arisen through multiple independent transmissions of simian immunodeficiency viruses from non-human primates. However, among these zoonotic events, only that leading to the HIV-1 group M lineage has achieved pandemic human-to-human transmission, infecting >80 million individuals(D’Arc et al., 2015; Gao et al., 1999; Gao et al., 1992). Other HIV lineages have led to lower numbers of detected infections: HIV-1(N) =<20, HIV-1(P)=2, HIV-2(F)=2 and HIV-2(C), (D), (E) (H)=1 each(Mourez et al., 2013; Plantier et al., 2009; Simon et al., 1998; Vallari et al., 2011). HIV-1(O) has infected around 100,00 humans(Mourez et al., 2013; Vessiere et al., 2010), while HIV-2(A) and (B) have collectively infected around 2 million individuals and have declined in number (De Cock et al., 1993; Gottlieb et al., 2008). It is unclear whether special features of HIV-1(M) have facilitated pandemic spread. Previous work has suggested that chance events have impacted development of the HIV pandemic(Faria et al., 2014) and a central role for HIV-1(M) Vpu adaptation to antagonise human tetherin has been proposed(Gupta and Towers, 2009; Sauter et al., 2009). Further, the cytoplasmic sensor cyclic GMP-AMP synthase (cGAS) has been shown to inhibit HIV-2 infectivity with a role for the nuclear protein Non-POU domain-containing octamer-binding protein (NONO) targeting the HIV-2 capsid(Lahaye et al., 2018). We hypothesised that specific HIV-1(M) adaptation in capsid might favour human-to-human transmission through evasion of innate immune sensing. The HIV core accommodates and regulates reverse transcription of the viral RNA genome into DNA(Jacques et al., 2016), initiated in the cytoplasm and completed in the nucleus(Burdick et al., 2020). Current models suggest that if exposed in the cytoplasm, lentiviral DNA can be recognised by cGAS(Gao et al., 2013; Lahaye et al., 2013), which when stimulated, synthesises cyclic guanosine monophosphate-adenosine monophosphate (cGAMP), which activates stimulator of interferon genes (STING) to recruit tank binding kinase 1 (TBK1), leading to phosphorylation and nuclear import of transcription factor (interferon regulatory factor 3 (IRF3) (Ablasser et al., 2013; Liu et al., 2015; Sun et al., 2013; Zhang et al., 2019). IRF3 induces type-I interferon (IFN) expression and subsequent expression of interferon-stimulated genes (ISGs). The capsid itself can also act as a pathogen associated molecular pattern (PAMP) recognised by the restriction factor tripartite motif protein 5 (TRIM5) (Pertel et al., 2011). Human TRIM5 is not thought to restrict HIV-1(M) because cyclophilin A (CypA) recruitment to the incoming core shields HIV-1(M) from human TRIM5 restriction(Kim et al., 2019; Towers et al., 2003). However, species variants of TRIM5, for example the rhesus macaque TRIM5, forms a restrictive cage around HIV-1(M) cores, inhibiting viral uncoating and nuclear entry(Ganser-Pornillos et al., 2011; Skorupka et al., 2019; Stremlau et al., 2004). Coordination of TRIM5 trimers at the vertices of the hexameric TRIM5 cage, also drives TRIM5 mediated K63 linked ubiquitin (Ub) chain synthesis, activation of TGFβ activated kinase 1 (TAK1) and subsequently AP-1 and NF-kB transcription factors(Fletcher et al., 2015; Fletcher et al., 2018; Pertel et al., 2011). Both cGAS and TRIM5 drive pro-inflammatory signalling responses expected to supress viral replication(Gao et al., 2013; Pertel et al., 2011). We hypothesised that pandemic HIV-1(M) should be particularly effective in avoiding sensing because early inflammatory responses, and interferon production, are expected to limit lentiviral transmission(Sandler et al., 2014). Here, we demonstrate that distinct pro-inflammatory profiles are induced following activation of cGAS (CXCL10, IFIT1, CCL5, MxA, IFIT2) and TRIM5 (IL8, IL1β, PTGS2, SOD2) by non-pandemic HIV-2 and HIV-1(O), but not pandemic HIV-1(M). We demonstrate unique structural adaptations in HIV-1(M) capsid that link to cGAS and TRIM5 insensitivity and propose that non-pandemic HIV strains lack the evolutionary adaptations required for effective innate immune evasion, limiting their human-to-human transmission and pandemic potential.

## RESULTS

### Non-pandemic HIV activate innate immunity

Pandemic HIV-1(M) isolates efficiently infect macrophages *in vitro*, in mouse models, and *in vivo*(Ganor et al., 2019; Honeycutt et al., 2016; Jambo et al., 2014; Rasaiyaah et al., 2013). Conversely, HIV-2 fails to replicate in macrophages and dendritic cells due in part to enhanced viral DNA synthesis, driven by Vpx mediated degradation of SAMHD1, sensing of viral DNA by cGAS and consequent innate immune activation(Chauveau et al., 2015; Duvall et al., 2007; Lahaye et al., 2013). We found that both Vpx-encoding HIV-2, and HIV-1(O) which doesn’t encode Vpx, cannot replicate in primary human monocyte-derived macrophages (MDM) unless replication was rescued by suppression of type-I interferon (IFN) signalling with an IFN receptor (IFNAR1) antibody (Ab) (Figure 1A-C). HIV-1 (M) replicated well and IFNAR1-Ab had no effect (Figure 1A)(Rasaiyaah et al., 2013). IFNAR1-Ab also enhanced single round infection in MDM with full-length HIV-2, or HIV-1(O), but not HIV-1(M) (Figure S1A). All viruses replicated efficiently in the permissive cell line GHOST demonstrating replication fitness (Figure S1B). Concordantly, infection of MDM with equal genome copies (qRT-PCR) of VSV-G pseudotyped HIV-2 and HIV-1(O) GFP encoding vectors, hereafter referred to as HIV-GFP, induced expression of ISGs (CCL5, IFIT1, MxA, CXCL10) and pro-inflammatory genes (IL-8, IL-1β, PTGS2 and SOD2), with IL-8 and CXCL10 secretion evidenced by ELISA (Figure 1D-F). Despite similar infectivity, equal genome copies of HIV-1(M) induced less ISG and cytokine expression, consistent with replication in MDM that was independent of interferon suppression (Figure 1A, D-F). DNA transfection and LPS treatment acted as positive controls for PAMP induction of innate responses (Figure 1E, F).

**Figure 1.**
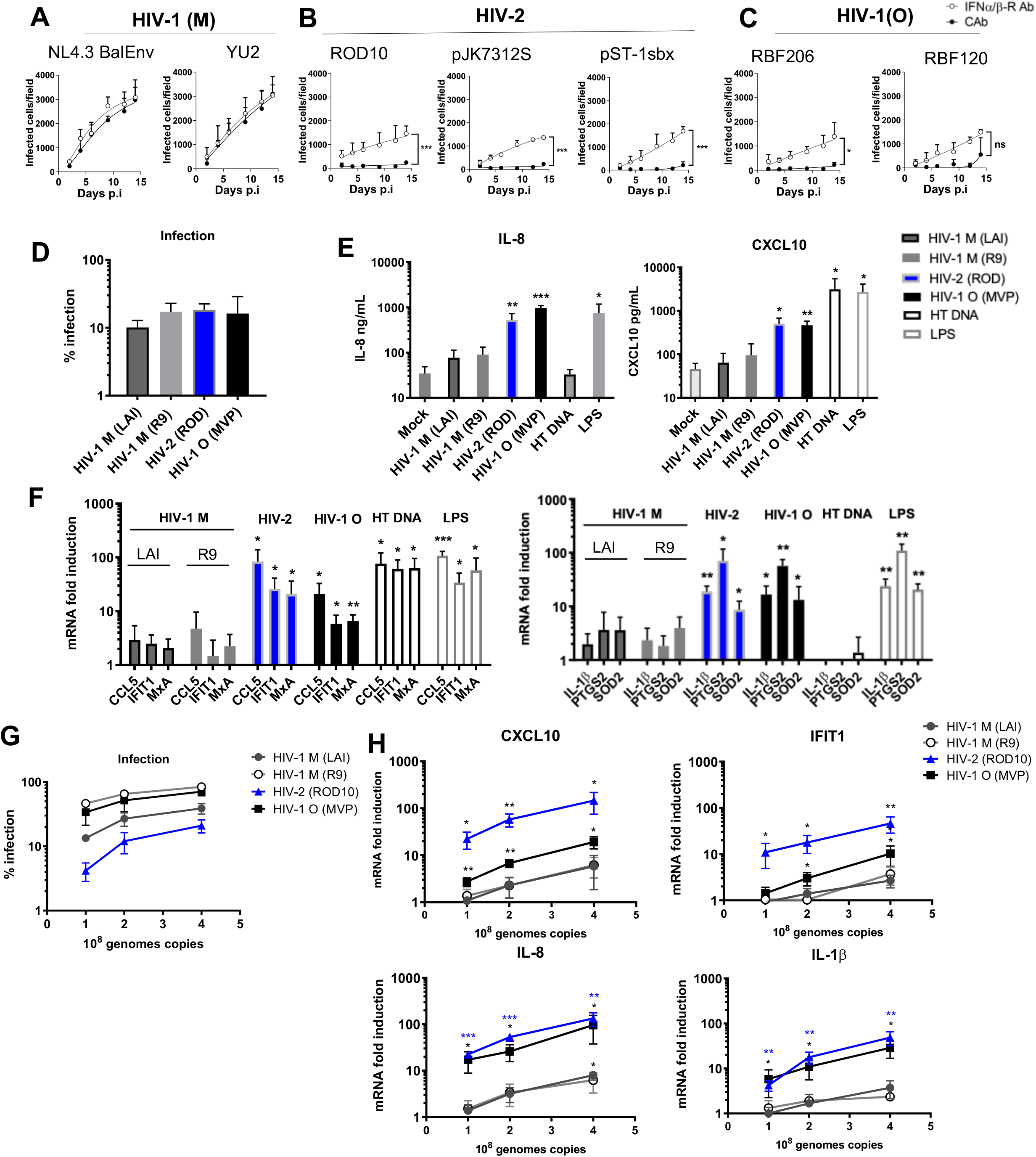
Non-pandemic HIV activate innate immune responses in macrophages more strongly than pandemic HIV-1(M). **(A)** Replication of HIV-1(M) or **(B)** HIV-2 or **(C)** HIV-1(O) isolates in human monocyte-derived macrophages (MDM) in the presence of interferon α/β receptor (IFNα/β-R) or control antibody (cAb). **(D)** Single round infection of MDM with equal genome copies of VSV-G pseudotyped HIV-1(M), HIV-2 and HIV-1(O) -GFP measured 48h post-infection. **(E)** Secreted IL-8 and CXCL10 from infections in (D) measured by ELISA 48h post-infection. (F) GAPDH-normalised mRNA levels in infections from (D) expressed as fold induction over untreated MDM 24h post-infection or after Herring testes (HT) DNA transfection (1ug/mL) or LPS stimulation (100ng/mL) (G) Infection of THP-1 cells with equal genome copies of VSV-G pseudotyped HIV-1(M), HIV-2 and HIV-1(O) –GFP measured 48h post-infection. (H) GAPDH-normalised mRNA levels from infections in (G) expressed as fold induction over untreated THP-1 cells 24h post-infection. Mean +/− SD, N=3 donors (A-E) or independent experiments (F-H). A-C, 2-way ANOVA vs Ctrl Ab. D, E, F unpaired t-test vs untreated MDM. G, H paired t-test vs untreated THP-1 cells. *p<0.05, **p<0.01, ***p<0.001

The striking difference in innate immune activation between pandemic and non-pandemic viruses was reproduced on infection of undifferentiated THP-1 cells, a myeloid cell line model (Figure 1G-H) (Tomlinson et al., 2012). Using VSV-G pseudotyped GFP encoding vectors, and equalising dose by genome copies (qRT-PCR), we found that HIV-2 and HIV-1(O) induced dose-dependent ISG and cytokine expression more strongly than HIV-1(M), even though HIV-1(M) infection levels were higher, ruling out higher infection levels as an explanation for enhanced gene induction by non-pandemic vectors (Figure 1G, H). Additionally, THP-1 bearing an ISG54 minimal promoter in conjunction with five IFN-stimulated response elements (IRF), or NF-kB-sensitive reporters were activated more strongly by HIV-2 and HIV-1(O), than HIV-1(M) infection (Figure S1C, D). Importantly, measurement of viral DNA synthesis, after infection with equal genome copies, revealed higher levels of infection, but similar levels of reverse transcripts, for HIV-1(M), also ruling out enhanced DNA synthesis by HIV-2/HIV-1(O) in THP-1 as the reason for increased innate activation (Figure S1E, F, G). HIV-1(M) is more efficient in infecting THP-1 cells per reverse transcript (Figure S1E, F, G) suggesting that extra non-infectious HIV-2/HIV-1(O) DNA may contribute to activating the immune response. RT products are plotted per infected cell to illustrate these differences in efficiency of infection with respect to DNA synthesis illustrating that HIV-1(M) has close to one infectious unit per molecule of DNA synthesised in these cells (Figure S1H). Importantly, VSV-G pseudotyped near full length HIV-1(M) LAI ΔEnv(Peden et al., 1991) (labelled LAI) and minimal HIV-1(M) R9 based GFP encoding vector (p8.91+CSGW(Bainbridge et al., 2001; Zufferey et al., 1997), labelled R9), gave similar low levels of stimulation in MDM and THP-1 cells (Figure 1E-H).

### cGAS and TRIM5 detect non-pandemic HIV

We hypothesised that non-pandemic innate immune activation reflected enhanced sensitivity to host cell sensing. Indeed, RNAi mediated depletion of well characterised HIV sensors cGAS and TRIM5 (Figure 2A) reduced induction of cGAS and TRIM5 specific genes by HIV-2 and HIV-1(O) in MDM. For example, induction of IFIT1, IFIT2 and CXCL10, by HIV-2 and HIV-1(O) was reduced by cGAS depletion (Figure, 2B) but not TRIM5 depletion (Figure 2C). Conversely, induction of IL-1β, PTGS2, and IL-8, were reduced by TRIM5 depletion (Figure 2E), but not cGAS depletion (Figure 2D). We also note that PTGS2, IL-1β and IL-8 were also not induced by transfecting DNA, consistent with them not being induced by cGAS activation (Figure 1E, F), As before, pandemic HIV-1(M) induced cGAS and TRIM5 sensitive genes poorly (Figure 2B-E).

**Figure 2.**
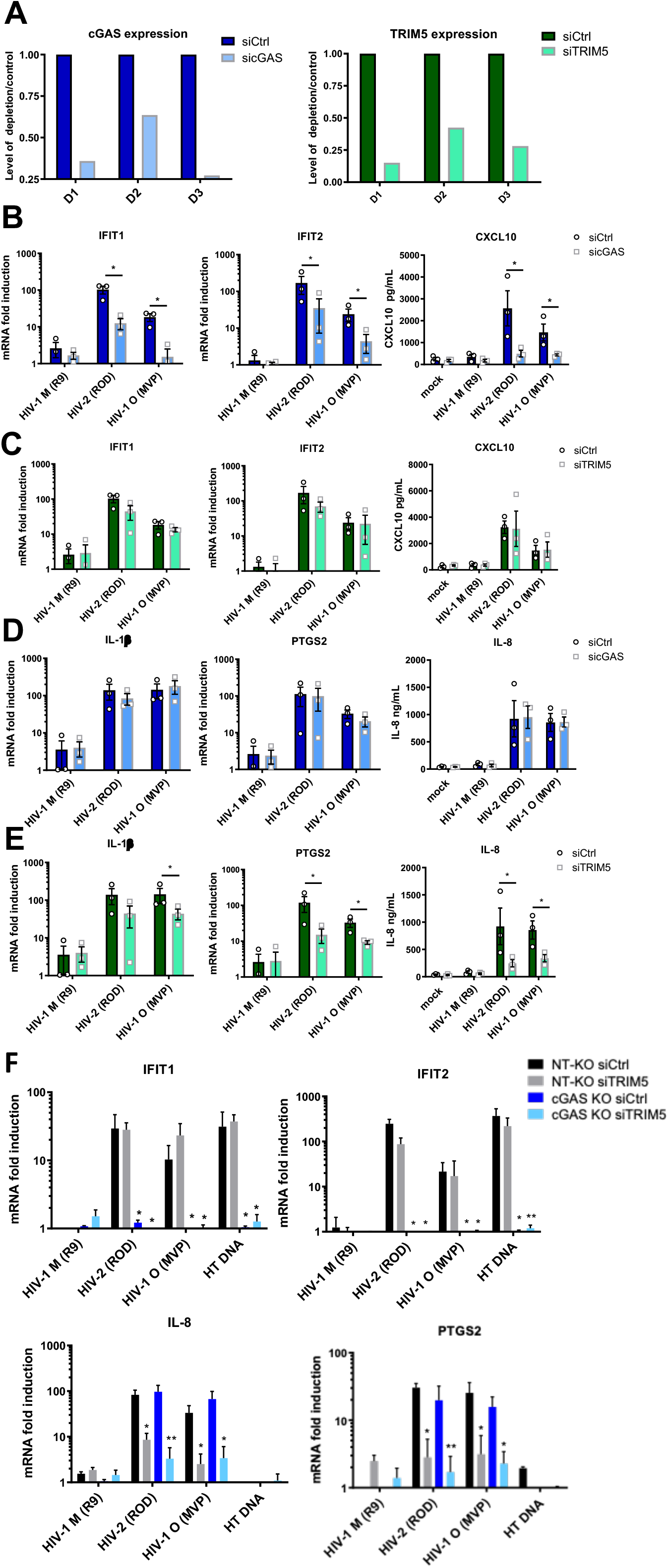
cGAS and TRIM5 detect non-pandemic HIV. **(A)** Example of GAPDH-normalised cGAS mRNA levels in MDM representing 3 independent donors transfected with cGAS targeting siRNA (sicGAS), TRIM5 targeting siRNA (siTRIM5) or non-targeting control (siCtrl). **(B-E)** GAPDH-normalised mRNA levels, or secreted cytokine levels (ELISA) (CXCL10, IL-8), expressed as fold induction over uninfected (mock) 24h post-infection (mRNA) or 48h post-infection (cytokine) from cells in (A). **(F)** GAPDH-normalised mRNA levels, expressed as fold induction over uninfected samples, in non-targeting CRISPR treated cells (NT-KO) transfected with control siRNA (siCtrl) (NT-KO siCtrl), NT-KO siTRIM5, cGAS KO siCtrl or cGAS KO siTRIM5 THP-1 cells 24h post-infection or after HT-DNA (1ug/mL) transfection. Mean +/− SD, n=3 independent experiments and donors. (B-E) unpaired t-test vs untreated MDM. (F) Paired t-test vs THP-1 Ctrl vector.*p<0.05, **p<0.01

Sensor specific gene induction by HIV-2 and HIV-1(O) was confirmed in THP-1 cells. We obtained commercial CRISPR/Cas9 ablated cGAS knock out THP-1 lines (Invivogen) and used them to show that cGAS knock out abrogated cGAS-sensitive IFIT1 and IFIT2 induction, but did not particularly impact induction of TRIM5-sensitive IL-8 and PTGS2 (Figure 2F dark blue bars). By contrast, siRNA mediated depletion of TRIM5 in THP-1 (Figure S2A) reduced TRIM5-sensitive IL-8 and PTGS2 induction, but had little effect on cGAS-sensitive IFIT1 and IFIT2 expression (Figure 2F, grey bars). Importantly, depleting TRIM5 with siRNA in the cGAS KO line (Figure S2A) reduced induction of all genes (Figure 2F, light blue bars). Knockout of RNA sensing adaptor MAVS had no impact on innate immune activation by HIV-2/HIV-1 (O) (Figure S2B-E). These data suggest that cGAS and TRIM5 independently contribute to the inflammatory innate immune response to non-pandemic lentivirus infection in macrophages.

### Nuclease TREX1 suppresses innate immune sensing of non-pandemic HIV

We expected TRIM5 to sense viral capsid and cGAS to sense viral DNA. To probe this we first used TREX1. Depletion of cytoplasmic nuclease TREX1 has been shown to cause HIV-1(M) DNA to activate cGAS(Towers and Noursadeghi, 2014; Yan et al., 2010), suggesting that DNA released from prematurely uncoating viral particles can trigger cGAS, if not degraded by TREX1. We therefore tested whether TREX1 over-expression in THP-1 cells could degrade the cGAS-activating viral DNA from non-pandemic viruses and suppress sensing. In fact, 20-fold TREX1 mRNA over-expression (Figure S3A), reduced ISG induction by HIV-2 and HIV-1(O), consistent with exposure of viral DNA activating innate immune response against both non-pandemic HIV (Figure 3A, B). Unexpectedly, TREX1 overexpression also reduced HIV-2 infectivity, with no effect on HIV-1(M) (Figure 3C) suggesting that HIV-2 capsids do not effectively shield viral DNA from high TREX1 levels. In contrast, HIV-1(O) infection was not impacted by TREX1 over-expression (Figure 3D), despite cGAS activation (Figure 1, 2B, D). This suggests that TREX1 sensitive HIV-2 is infectious and that HIV-2 and HIV-1(O) are different in their uncoating strategies, perhaps regarding timing and location, leading to different levels of TREX1 sensitivity and cGAS activation.

**Figure 3.**
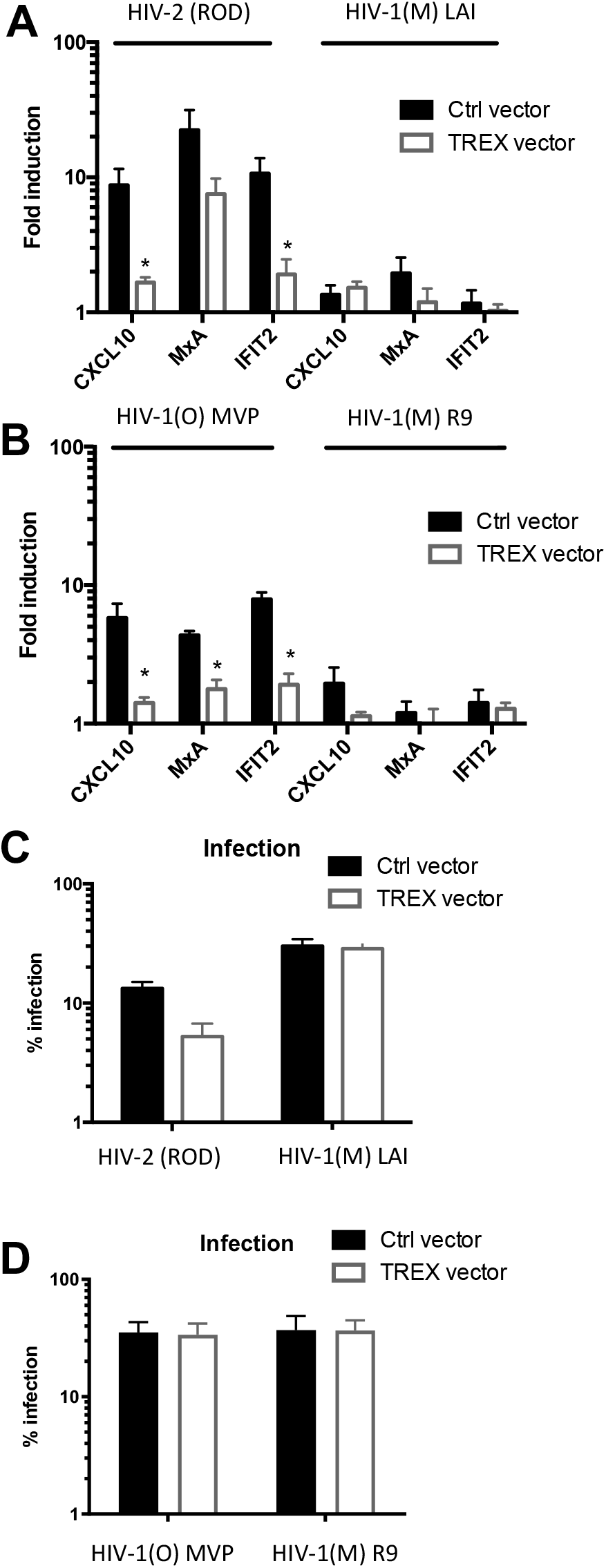
Nuclease TREX1 suppresses innate immune sensing of non pandemic HIV. **(A, B)** GAPDH-normalised ISG mRNA levels, expressed as fold induction over uninfected, in control vector (Ctrl) or TREX1-expressing THP-1 cells with HIV-2 and HIV-1(M) or HIV-1(O) with infection levels in parallel wells measured 48 hpi shown in **(C, D)**. Mean+/− SD, n= 3. B, D paired t-test vs THP-1 Ctrl vector. *p<0.05

### Genome free HIV particles activate TRIM5, but not cGAS, and trigger an antiviral state

To assess the contribution of TRIM5 capsid-sensing to host gene induction we used VSV-G pseudotyped viral-like particles (VLP), produced without genomes. As expected, genome-free VLP induced TRIM5-sensitive IL-8, IL-1β, PTGS2, but not cGAS-sensitive IFIT1/IFIT2 in THP-1 (Figure 4A). To test whether TRIM5 activation induced an antiviral state, we activated TRIM5 in THP-1 using increasing doses of genome free VLP, and then infected the same cells 24h later. As a control for innate activation, we observed dose dependent activation of TRIM5 sensitive NF-kB reporter by HIV-2 and HIV-(O) VLPs, but very little activation by HIV-1(M) VLPs (Figure S4A). Importantly, responses to non-pandemic VLP inhibited a second infection by HIV-2 or HIV-1(O)-GFP (Figure 4B). Consistent with the failure of HIV-1(M) to activate gene expression, HIV-1(M) VLPs did not induce inhibition of HIV-2 or HIV-1(O)-GFP infection (Figure 4B). Surprisingly, HIV-1(M) was also insensitive to the effects of prior exposure to HIV-2 or HIV-1(O) VLPs (Figure 4B). Thus, TRIM5 activation induces the expression of antiviral genes, including secretion of IL-1β (Figure S4B) that can restrict non-pandemic HIV. Concordantly, pre-treatment of THP-1 with IL-1β, or IFN-β, reduced infection of pandemic and non-pandemic HIV strains, but pandemic HIV-1(M) were notably less sensitive to IFN-β, and insensitive to IL-1β even when the target cells were pretreated (Figure S4C, D). Importantly, single round infections with all viruses were much less sensitive to IL-1β and IFN treatment when added, for example, 6 hours post-infection (Figure S4D). This is consistent with a model in which single round infections induce cytokines too late to inhibit that first round of infection. This is also evidenced by modest rescue of single round infection (GFP expression) of HIV-2 and HIV-1(O) when TRIM5 is depleted in macrophages (Figure S4E). We propose that sensor activation initiates a response that has stronger effects on later rounds of infection, as evidenced by the inhibition of infection when cells are pre-treated with IFN or IL-1β (Figure S4D).

**Figure 4.**
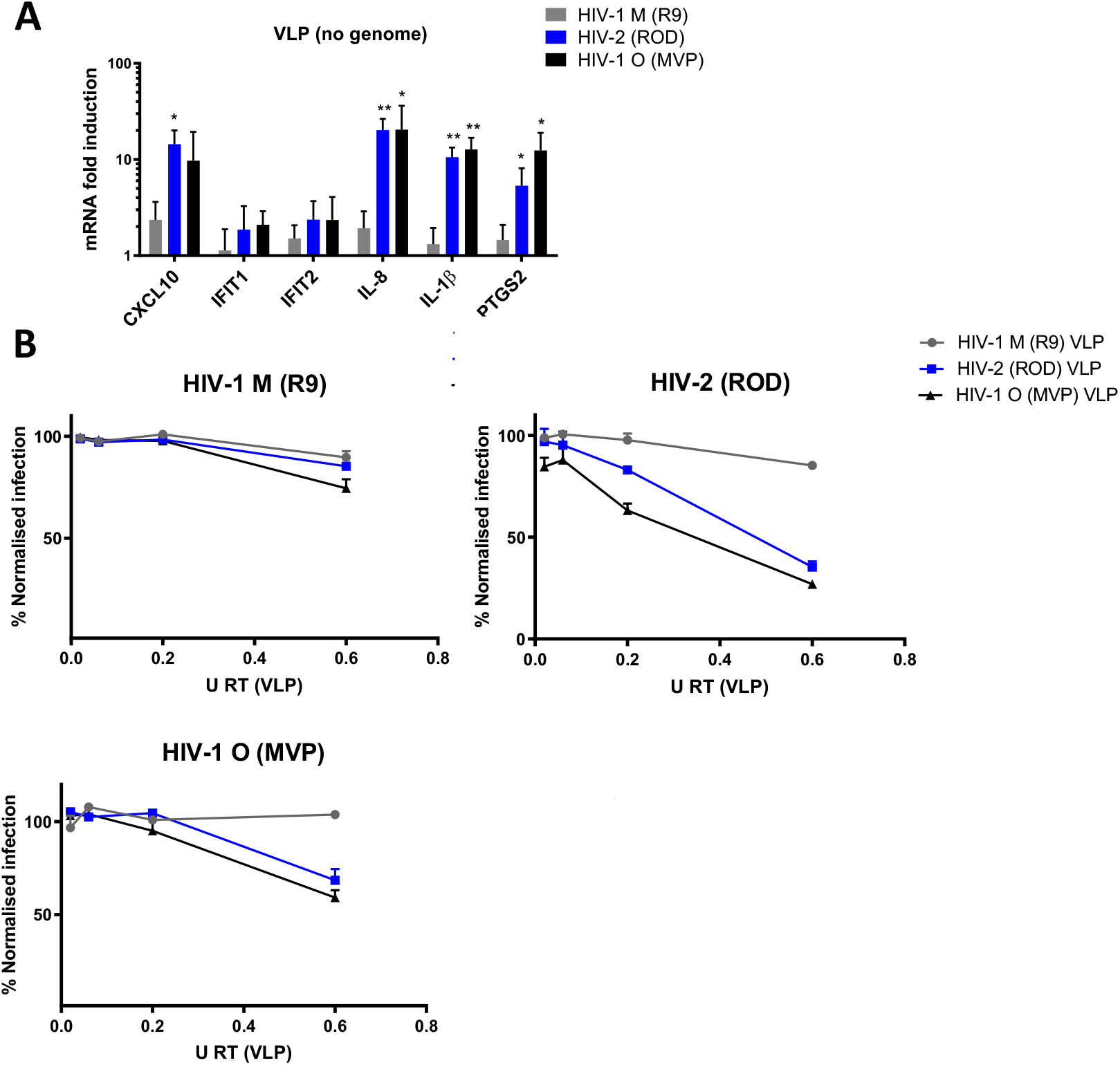
Genome free HIV particles activate TRIM5, but not cGAS, and trigger an antiviral state. **(A)** GAPDH-normalised mRNA levels expressed as fold induction over uninfected THP-1 cells 24h post viral-like particle (VLP) treatment. **(B)** HIV-1(M), HIV-2 and HIV-1(O) infection levels normalised to levels of infection without prior exposure to VLP. Mean +/− SD, n=3 (A) Paired t-test vs untreated THP-1 *p<0.05, **p<0.01. VSV-G pseudotyped VLP are made without genome.

### Lineage associated adaptation of HIV capsid at position 50

Lentiviral capsids protect viral nucleic acids from detection by sensors and are the target for TRIM5 restriction(Fletcher et al., 2018; Siddiqui et al., 2019; Stremlau et al., 2004; Sumner et al., 2020). Indeed, an important role for capsid in determining interferon induction in macrophages was confirmed by our observation that HIV-1(M) chimeras bearing HIV-1(O) or HIV-2 CA could only replicate in MDM in the presence of IFNAR1-Ab, (Figure 5A). Thus the chimeric viruses bearing HIV-1(O) or HIV-2 capsid proteins behaved similarly to wild type (WT) HIV-2/HIV-1(O) in these cells (Figure 1A-C). Importantly, the chimeric viruses replicated efficiently in GHOST cells indicating replication fitness (Figure S5A). Previously reported X-ray structures of HIV-1(M) CA hexamers suggest that they can crystalise in two distinct conformations, with respect to a channel at the 6-fold symmetry axis. The position adopted by the beta hairpin structure (BHP) above the channel can be open (His12 forms a salt bridge with Asp51) or closed (in which there is a bridging water molecule between His 12 and Asp 51) (Figure 5B)(Jacques et al., 2016). Within the channel, six arginines (R18) form a positively charged centre suggesting a model in which nucleotides are electrostatically recruited into HIV cores to fuel encapsidated DNA synthesis(Jacques et al., 2016; Xu et al., 2020). We proposed that this mechanism is conserved in HIV-1(O) and HIV-2 because they conserve Arg at the equivalent positions at the 6-fold symmetry axis(Jacques et al., 2016). Furthermore, like HIV-1(M), purified recombinant HIV-1(O) hexamer can bind dCTP at nanomolar Kd measured by fluorescence anisotropy (Figure S5B). A role for channel associated arginines in HIV-1(O) and HIV-2 DNA synthesis was further supported by titrating mutant packaging constructs bearing arginine-to-glycine substitutions into HIV-2 GFP (R17G) and HIV-1(O) (R18G) GFP vectors to make chimeric viral particles with varying proportions of WT and R-to-G mutant CA protein. Increasing mutant CA proportionally reduced viral infectivity and DNA synthesis (Figure S5C). Indeed, DNA and infection profiles of WT/mutant mixtures of HIV-1(O) and HIV-2 were similar to those produced by titrating the HIV-1 R18G mutant into HIV-1(M) vector production transfections (Jacques et al., 2016).

**Figure 5.**
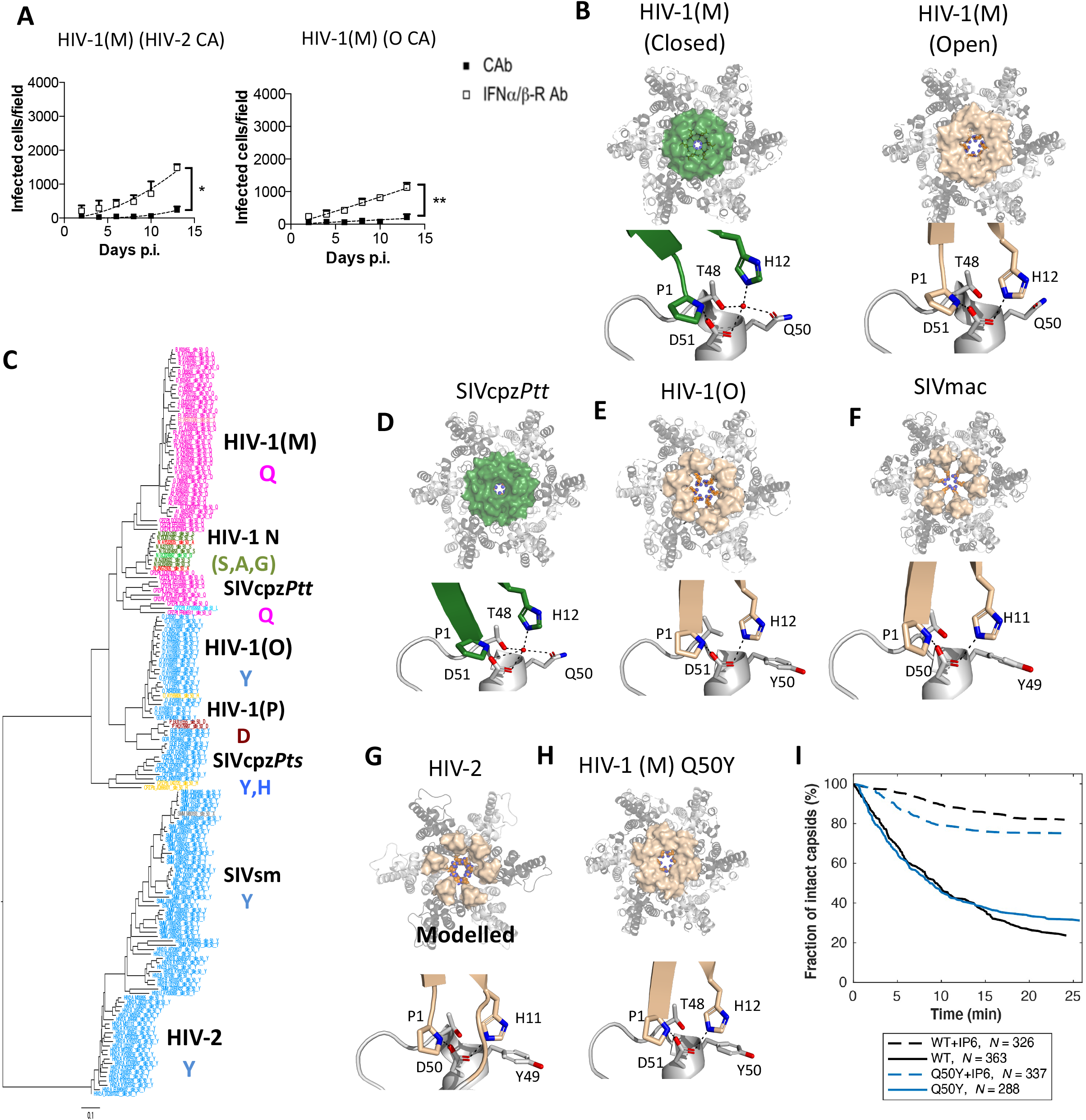
Pandemic associated adaptation of HIV capsid at position 50. **(A)** Replication of HIV-1(M) NL4.3 (BalEnv) bearing HIV-2 ROD10 or O MVP5180 Capsid in MDM in the presence of anti-interferon α/β receptor (IFNα/β-R), or control, antibody (cAb). N=3 donors. A 2-way ANOVA vs Ctrl Ab. \**p*<0.05, \*\**p*<0.01, **(B)** HIV-1(M) CA hexamer highlighting β-hairpin (BHP) position in a closed (green) (PDB ID:5HGN) or open conformation (wheat) (PDB ID:5HGL). Lower panel details residues in hinge region that close BHP by coordinating water and increasing distance between His12 and Asp51. **(C)** Maximum likelihood phylogenetic tree of primate lentiviral capsid genes coloured by chromaclade to illustrate the residues equivalent to CA Q50 in HIV-1(M). **(D)** SIVcpz CA hexamer (PDB ID:7T15) with the β-hairpin (BHP) position in a closed (green) conformation. Lower panel details residues in hinge region that close BHP by coordinating water and increasing distance between His12 and Asp51. **(E, F)** HIV-1(O) (PDB ID: 7T12) and SIVmac (PDB ID:7T14) hexamers with open BHP (wheat) and key Gln at position 50 substituted for Tyr (Y) preventing water coordination and channel closure. **(G)** A modelled hexamer built from the HIV-2 N-terminal CA domain (PDB ID:2X82) is shown for comparison with HIV-1(M) and (O). The HIV-2 BHP (wheat) is open. Note HIV-2 position 49 is Tyr (Y). **(H)** Crystal structure of HIV-1(M) Q50Y (PDB ID:7T13) highlighting β-hairpin (BHP) position in an open conformation (wheat). R18 is shown at the centre of the hexamers. **(I)** Wild type and CA Q50Y capsid survival curves obtained from TIRF *in vitro* uncoating experiments. In the absence of IP6, HIV-1 Q50Y capsids are metastable and disassemble spontaneously with similar half life to wild type. Most capsids are stabilised in the presence of 100 μM IP6. Survival curves were generated from single-virion uncoating traces (*N*=326 for WT, *N*=326 for WT+100 μM IP6, *N*=326 for Q50Y, *N*=326 for Q50Y+100 μM IP6) from one representative uncoating experiment; see Figure S5E for additional Q50Y data.

To identify functional differences between HIV-1(M) and HIV-1(O)/HIV-2 capsids, we combined structural and evolutionary approaches. We first used Chromaclade to colour maximum likelihood phylogenetic capsid trees to highlight lineage defining amino acids(Monit et al., 2019). In this analysis the same phylogenetic tree is coloured according to the amino acid present at each position in the alignment. The coloured tree for CA position 50 (Figure 5C) stood out because the amino acid at this position defines the pandemic lineage, with HIV-1(M) and its chimpanzee parent (SIVcpz*Ptt*), uniquely bearing glutamine (Q) at CA50. SIV from Red Capped Mangabeys (SIVrcm), the parent providing capsid to SIVcpz*Ptt* (Bailes et al., 2003), SIVgor and HIV-1(O) all bear tyrosine (Y) at CA position 50. This is consistent with tyrosine being the ancestral state and Y50Q occurring in chimpanzees (Figure 5C). Tyrosine is also present in SIVsmm and maintained in HIV-2 consistent with it being ancestral. We note that Y50Q requires two nucleotide changes (UAY to CAR) consistent with adaptive change. Interestingly, the two non-pandemic HIV-1(P) strains both encode CA50D (Plantier et al., 2009), while HIV-1 (N) isolates encode CA50S, (n=6), CA50A (n=2) or CA50G (n=1) and again require two nucleotide substitutions from the ancestral Q50. These examples suggest different evolutionary pathways for CA50 adaptation in the different SIV and HIV lineages.

Crystal structures of HIV-1(O) capsid hexamers revealed that, unlike HIV-1(M), the HIV-1(O) hexamers exclusively adopted an open conformation (Figure 5E). Comparison of HIV-1(M) and HIV-1(O) hexamer structures explain the importance of position 50, which is located in the BHP hinge region (compare Figure 5B to 5E). In the HIV-1(M) hexamer, residue Q50, contributes to the tetrahedral hydrogen bonding network that promotes the closed BHP position. This is also true in the SIVcpz*Ptt* capsid that is parental to the pandemic HIV-1(M). An SIVcpz*Ptt* hexamer structure revealed that like HIV-1(M), Q50 contributes to the closed BHP position by coordinating a water molecule (Figure 5D). However, in HIV-1(O) Y50 has been retained from the ancestral SIVgor, preventing water coordination and BHP closure (Figure 5E). All known HIV-2 and SIVsmm CA sequences bear tyrosine at this position consistent with a conserved structural difference to pandemic HIV-1 (Figure 5C). We have been unable to crystalise the HIV-2 CA hexamer. However, comparison of the HIV-1(O) hexamer hinge region with the hinge region of the published HIV-2 CA N-terminal domain structure (PDB ID 2X82) (Ylinen et al., 2010) and a modelled HIV-2 hexamer, illustrates the resemblance between these non-pandemic capsids, and suggests an open BHP position for HIV-2 due to the similarity in the region around Y49 in HIV-2 and Y50 in HIV-1(O) (Figure 5E, G). We were able to solve the structure of an SIVmac hexamer (Figure 5F). SIV from macaques (SIVmac) was unknowingly transmitted during laboratory experiments from Sooty Mangabeys infected with SIV Sooty Mangabey (SIVsmm) to Rhesus Macaques in captivity and is closely related to the parental SIVsmm virus (Figure 5C)(Apetrei et al., 2005). SIVmac hexamer retained the open BHP conformation and the structure of the BHP hinge region including the position Y50, which is highly conserved between HIV-1(O), HIV-2 and SIVmac further illustrating the structural similarity between hexamers from viruses bearing tyrosine at CA position 50 despite the HIV-1(O) and SIVsmm lineages being highly divergent (Figure 5E-G). To probe the role of CA50 in HIV-1 infectivity we made HIV-1(M) CA Q50Y, reverting the glutamine at position 50 to the ancestral tyrosine. Solving the mutant hexamer structure confirmed that the BHP of HIV-1(M) CA Q50Y adopted an open conformation and the hinge region of the HIV-1(M) Q50Y mutant resembled the HIV-1(O) and SIVmac structures (Figure 5H). Indeed, overlay of the open WT HIV-1(M) hexamer structure (PDB ID: 5HGL) and HIV-1(M) CA Q50Y (PDB ID:7T13) demonstrated the similarity between them (Figure S5D). Importantly, and unlike other capsid point mutants for example CA K25A which prevents IP6 recruitment into assembling virions (Renner et al., 2021), HIV-1 (M) Q50Y capsids were not intrinsically destabilised, showing wild type disassembly kinetics in the absence of IP6 in single molecule uncoating assays (Figure 5I and S5E) (Marquez et al., 2018). As expected, given the similar structure, addition of IP6 led to an increase of the HIV-1(M) Q50Y capsid half life from minutes to hours for the majority of capsids, confirming that IP6 binding was not perturbed. Nevertheless, in infection experiments, single round infectivity of HIV-1(M) GFP CA Q50Y in MDM was poor and this capsid mutant did not replicate (Figure S5F).

### Adaptation of HIV capsid at position 120

Given that HIV-1 CA Q50Y was defective we returned to Chromaclade (Monit et al., 2019) seeking additional changes, with some lineage specificity, that occurred along with capsid Y50Q in SIVcpz*Ptt*. This revealed a patch of differences around CA position 120, identified as a deletion of Arginine CA 120 in HIV-1(M) and its SIVcpz*Ptt* parent (Figure 6A). Inspection of the non-pandemic hexamer structures revealed that the arginine lost from SIVcpz*Ptt*/HIV-1(M) forms a salt bridge between helix six and the CypA binding loop in HIV-1(O) and HIV-2 CA helix 6: Arg 120-Glu 98 in HIV-1(O), Arg 118-Glu 96 in HIV-2 (Figure 6B). This salt bridge is also conserved in the SIVmac hexamer, between Arg 117 and Glu 95, but not in the SIVcpz*Ptt* structure (Figure 6B). These observations suggest that, in addition to Y50Q, the SIVcpz*Ptt* parent of the pandemic HIV-1(M) lineage lost a salt bridge on the CA surface through loss of an arginine residue. To examine its phenotypic effect, we reversed the deletion in HIV-1(M), restoring the arginine in mutant HIV-1(M)+R120. The X-ray structure of this HIV-1(M) CA +R120 hexamer mutant revealed that adding the arginine reinstated the salt bridge (Figure 6B) and this restored the infectivity of VSV-G pseudotyped HIV-1(M) CA Q50Y+R120 in MDM (Figure 6C)

**Figure 6.**
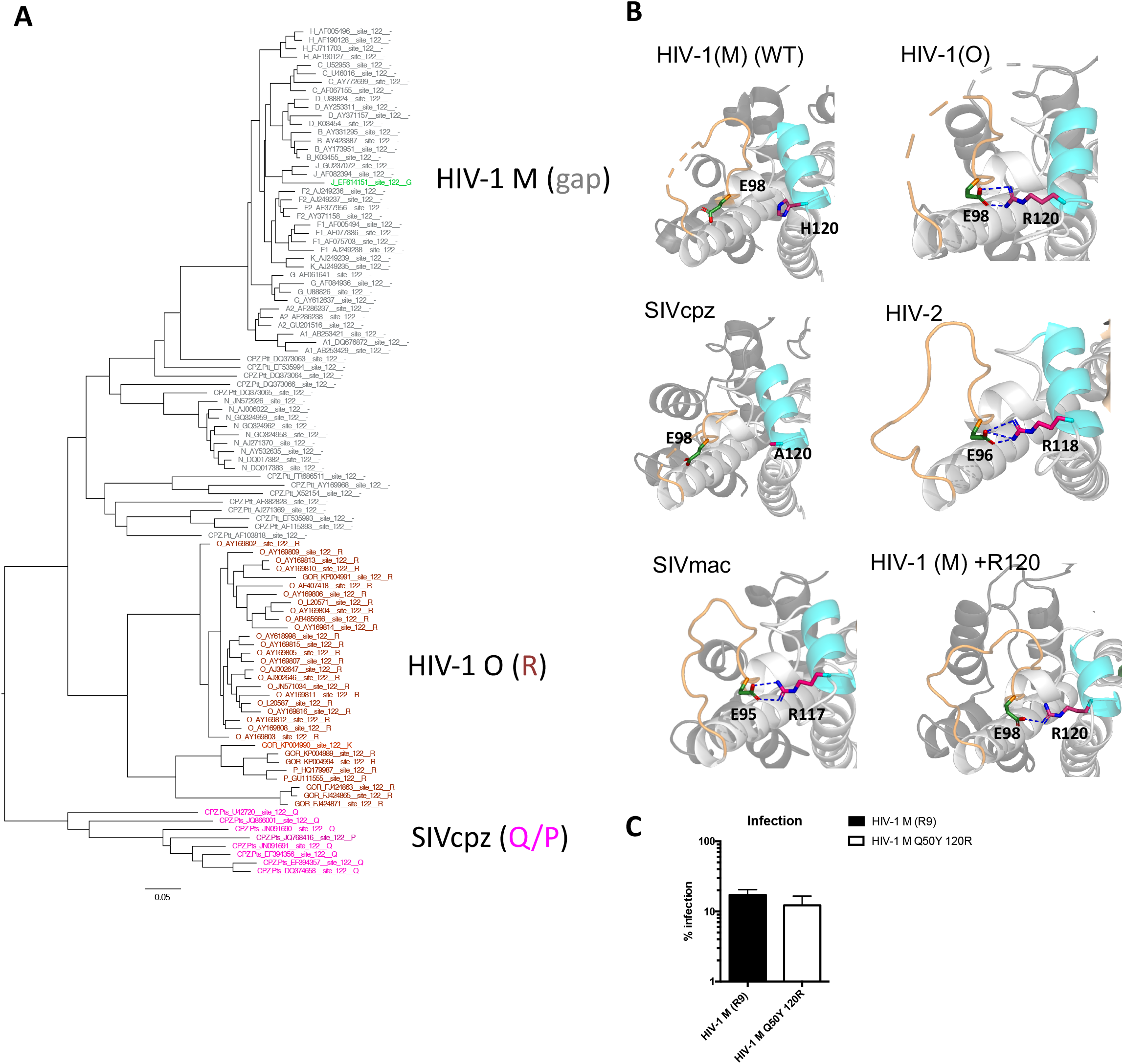
Pandemic associated adaptation of HIV capsid at position 120. **(A)** Maximum likelihood phylogenetic tree of primate lentiviral capsid genes coloured to illustrate the residues equivalent to HIV-1(O) CA 120. Grey and (–) branch labels denotes a gap in the alignment. **(B)** Structures showing salt bridges in HIV-1(O) (PDB ID:7T12) (E98-R120), HIV-2 (PDB ID:2X82) (E96-R118), SIVmac (PDB ID:7T14) (E95-R117) and HIV-1(M) +R120 (PDB ID:7QDF) (E98-R120). The salt bridge is absent in wild type (WT) HIV-1(M) CA (PDB ID:5HGN) and SIVcpz (PDB ID:7T15) because R120 is absent. **(C)** Single round infection of MDM with equal genome copies of VSV-G pseudotyped HIV-1(M) or HIV-1(M) CA Q50Y 120R –GFP measured 48h post infection by flow. Mean +/− SEM n=3.

### Capsid Q50Y 120R mutations makes pandemic HIV-1(M) sensitive to cGAS and TRIM5

We next investigated whether pandemic lineage-associated mutations influenced host responses to infection and macrophage tropism. To test this, we reversed the pandemic-associated CA mutations in HIV-1(M) and infected MDM. We found that HIV-1(M) CA Q50Y+R120 could only replicate in MDM in the presence of IFNAR1-Ab, consistent with enhanced sensing and innate immune activation (Figure 7A). Concordantly, VSV-G pseudotyped HIV-1(M) CA Q50Y+R120, induced cGAS and TRIM5 sensitive genes in MDM more strongly than WT HIV-1(M), (Figure S6A, B – compare with Figure 1 E-F). cGAS-sensitive gene induction by HIV-1(M) CA Q50Y+R120 was reduced by cGAS depletion in MDM, (Figure 7B) and cGAS knockout in THP-1 (Figure 7D). TRIM5-sensitive gene induction was suppressed by TRIM5 depletion in MDM and THP-1 (Figure 7C-D). Importantly, combination of cGAS knock out and TRIM5 depletion in THP-1 suppressed all gene induction by HIV-1(M) CA Q50Y+R120 infection (Figure 7D). HIV-1(M) CA Q50Y+R120 VLPs without genome, induced TRIM5-sensitive, but not cGAS-sensitive, genes in THP-1 (Figure 7E). MAVS depletion had no effect on innate immune responses elicited by HIV-1(M) CA Q50Y+R120 infection (Figure S6C, D). Finally, we found that HIV-1(M) CA Q50Y+R120 behaved like non-pandemic viruses HIV-1(O) and HIV-2, in that it was more sensitive to restriction by human TRIM5 than wild type HIV-1(M), evidenced by rescue of infection by TRIM5 depletion in U87 cells, using stable shRNA expression (Figure S6E). Infection with TRIM5 sensitive MLV-N and insensitive MLV-B acted as controls (Keckesova et al., 2006). Thus, an HIV-1(M) CA mutation bearing Q50Y+R120, to represent non-pandemic HIV lineages, behaved like non-pandemic HIV, activating innate immune gene expression in a TRIM5 and cGAS sensitive way. These observations outline how key amino acid changes in HIV-1(M) led to structural changes in the capsid that correspond with reduced activation of, or restriction by, key lentiviral sensors, cGAS and TRIM5. We therefore propose that the SIVcpz*Ptt*/HIV-1(M) lineage has undergone complex adaptations that underlie its pandemic success and that this study provides mechanistic insight into how particular pandemic lineage-specific adaptations at capsid positions 50 and 120 have paved the way for this particular SIVcpz*Ptt* lineage to become pandemic in humans as HIV-1(M).

**Figure 7.**
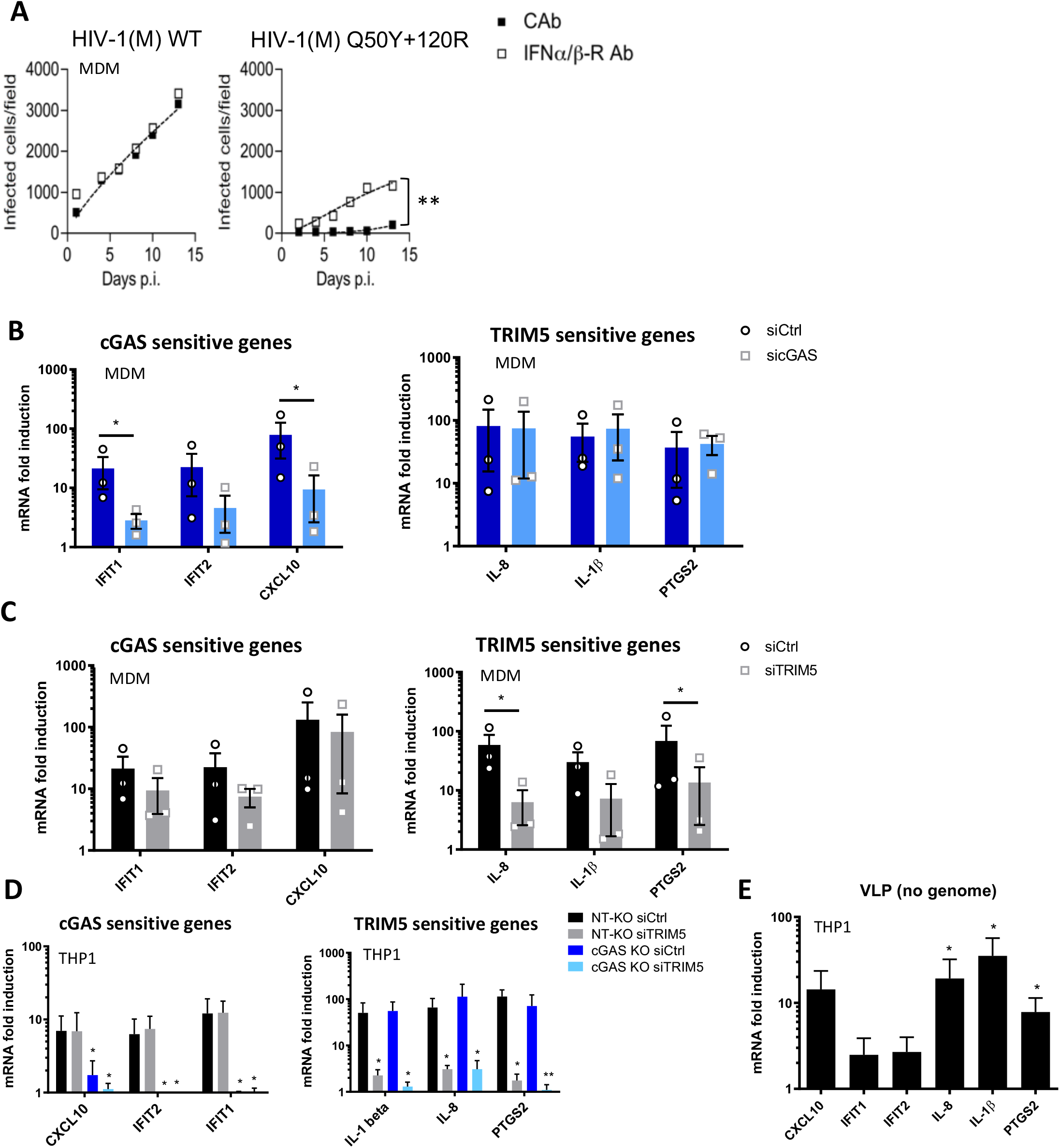
Capsid Q50Y +120R mutations make pandemic HIV-1(M) sensitive to cGAS and TRIM5 and insensitive to CypA. **(A)** Replication of HIV-1(M) NL4.3 (BalEnv) wild type (WT) or HIV-1(M) NL4.3 (BalEnv) bearing CA Q50Y +120R in MDM in the presence of interferon α/β receptor (IFNα/β-R) or control antibody (cAb). **(B)** GAPDH-normalised mRNA levels induced by HIV-1(M) CA Q50Y 120R, expressed as fold induction over uninfected samples in control siRNA transfected (siCtrl) or cGAS siRNA transfected (sicGAS) MDM 24h post-infection **(C)** GAPDH-normalised mRNA levels induced by HIV-1(M) CA Q50Y 120R expressed as fold induction over uninfected samples in control siRNA (siCtrl), or TRIM5 siRNA (siTRIM5), transfected MDM 24h post-infection. **(D)** GAPDH-normalised mRNA levels induced by HIV-1(M) CA Q50Y +120R expressed as fold induction over uninfected samples in non targeting CRISPR treated cells (NT-KO) transfected with control siRNA (siCtrl) (NT-KO siCtrl), NT-KO siTRIM5, cGAS KO siCtrl or cGAS KO siTRIM5 THP-1 cells measured 24h post-infection **(E)** GAPDH-normalised mRNA levels induced by HIV-1 (M) Q50Y +120R viral-like particles (VLP, no genome) expressed as fold induction over uninfected THP-1 cells measured 24h post-infection. Mean +/− SD, n= 3 independent experiments or donors. A. 2-way ANOVA vs Ctrl Ab. B, C unpaired t-test vs siCtrl. D, E paired t-test vs NT-KO siCtrl THP1 cells. \**p*<0.05, \*\**p*<0.01, \*\*\**p*<0.001.

## DISCUSSION

In this study, we set out to understand what is special about pandemic HIV-1 as compared to the other non-pandemic HIV lineages that are derived from independent zoonoses from simians. Accumulating data highlight the importance of HIV capsid in regulating viral DNA synthesis and shielding viral DNA from cytosolic sensors. Indeed, recent study of fluorescently labeled HIV-1 particles has shown that HIV-1(M) capsids remains intact during travel across the cytoplasm and through the nuclear pore complex, uncoating shortly before integration in the nucleus(Bejarano et al., 2019; Burdick et al., 2020; Li et al., 2021; Muller et al., 2021; Zila et al., 2021). Thus DNA synthesis takes place inside intact cores, during cytoplasmic transport, possibly completing in the nucleus, consistent with a key protective function for capsid against nucleases and innate immune sensors. Here, we have discovered that the SIVcpz*Ptt* parent of pandemic HIV-1(M) evolved specific adaptations in capsid that improve evasion of human innate immune sensors cGAS and TRIM5, as compared to non-pandemic strains derived from independent zoonoses.

We hypothesise that the structural differences that we observe between lentiviral capsids influence TRIM5 recruitment and subsequent inhibition of infection. We note that TRIM5 depletion rescued single round infection of sensitive viruses in MDM (Figure S4E) to a similar degree as IFN receptor blockade (Figure S1A). TRIM5 depletion also suppressed macrophage activation of gene expression by non-pandemic HIV-2 and HIV-1(O) infection. These data suggest important roles for both innate immune signaling, and physical caging of incoming capsids, for TRIM5 antiviral activity (Li et al., 2016). TRIM5 signaling is driven by TRIM5 RING trimerisation at the vertices of the hexameric TRIM5 lattice that forms on viral cores (Fletcher et al., 2015; Fletcher et al., 2018) and HIV-1(M) appears to escape sensitivity to human TRIM5 in part through CypA recruitment (Kim et al., 2019). It is also possible that capsid surface dynamics influence TRIM5 recruitment and therefore the gain or loss of the salt bridge on the capsid surface also impacts TRIM5 restriction and activation of TRIM5 signaling. A complete understanding of TRIM5 sensitivity determinants awaits atomic level structural definition of lentiviral capsid-TRIM5 interactions(Li et al., 2016; Skorupka et al., 2019). We hypothesise that capsid influences cGAS sensitivity indirectly, through regulating the timing and position of uncoating and genome release, and therefore exposure of viral DNA PAMP to cGAS. This model is supported by our observation that over-expression of TREX1 suppresses innate immune activation by both HIV-1(O) and HIV-2 and inhibits infectivity of HIV-2 suggesting that partially uncoated, and therefore TREX1 sensitive, HIV-2 cores are infectious (Figure 3).

Our study focuses on the phenotypes of a small number of non-pandemic HIV lineages, but our phylogenetic analyses suggest significant complexity in HIV capsid evolution. For example, it is striking that the SIVcpz*Ptt* lineage gave rise to both pandemic HIV-1(M), which retained Q50, and non-pandemic HIV-1(N), which only infected a small number of people, but completely replaced Q50 with S, A or G, each requiring multiple nucleotide changes (Figure 5). On the other hand, Y50 is retained in SIVsmm to HIV-2 and chimpanzee-to-gorilla and gorilla-to-human HIV-1(O) zoonoses (Figure 5). Furthermore, the SIVcpz*Ptt* progenitor also gave rise to viruses retaining the ancestral tyrosine at CA50 (SIVgor, HIV-1(O)) suggesting an unsampled lineage of SIVcpz*Ptt* that kept this amino acid. While we cannot be sure of the pressures that selected capsid adaptations, we hypothesise that complexity arises from the diversity of capsid function and the species-specific cofactor interactions that govern regulation of DNA synthesis, uncoating and nuclear entry which frequently, but not always require adaptation of the hexamer channel. Certainly there is more to pandemic adaptation than changes at capsid positions 50 and 120, because we were unable to enhance HIV-2/HIV-1(O) replication in human MDM by mutating CA Y50Q, which significantly reduced infectivity of both HIV-1(O) and HIV-2 (data not shown).

HIV-2 and HIV-1(O) replicate efficiently in activated primary T cells *in vitro*(Chauveau et al., 2015; Dittmar et al., 1999; Mack et al., 2017). Here we focused on primary human macrophage replication because primary T-cells are not permissive to HIV replication unless activated, typically by cross-linking the T-cell receptor (TCR) to mimic antigen stimulation. Thus, TCR-dependent signalling and cytokine secretion dominate *in vitro* HIV infection experiments, obviating experimental interpretation of virus-induced changes such as activation of cGAS/TRIM5 signaling. In addition to being permissive to infection, macrophages also effectively secrete type-I IFN which can inhibit lentiviral transmission in animals (Sandler et al., 2014) and is present during transmission-induced cytokine storms in humans (Stacey et al., 2009). An important role for IFN in HIV transmission is evidenced by the unique resistance of transmitted founder HIV-1 to type-I interferons (Iyer et al., 2017). Previous work has demonstrated the antagonistic role of type-I interferons and IL-1β, for example how IFN can downregulate IL-1β production and inflammasome-mediated IL-1β processing (Guarda et al., 2011; Hu et al., 2005; Mayer-Barber and Yan, 2017). However, recent studies have shown that IL-1β drives the induction of type-I IFN after dengue, zika or vesicular stomatitis virus infection (Aarreberg et al., 2019; Orzalli et al., 2018), contributing to the antiviral response. These observations highlight how the regulation of IFN and IL-1β over the course of a viral infection (Gondim et al., 2021) and how cytokine responses interact and influence each other to promote an antiviral state is still not well understood. Nontheless, together with our data, these observations emphasise the importance of innate immune evasion as a key determinant of transmission and therefore pandemic potential. We note that the changes that allow evasion of innate sensing occurred in viruses infecting central chimpanzees (*Pan troglodytes troglodytes*) prior to transmission to humans, suggesting that SIVcpz*Ptt* are more dangerous to humans than SIVcpzPts strains infecting eastern chimpanzees (*Pan troglodytes schweinfurthii*), which have never been detected in the human population. This in turn suggests that we may be able to predict zoonosis competent viruses by examining their capacity to antagonise or evade human innate immunity prior to human adaptation. We note that the first wave of SARS-CoV-2 in humans effectively antagonised human innate immunity, despite likely bat origins, but that the pandemic Alpha variant has enhanced these properties, again linking enhanced innate immune evasion to pandemic human-to-human spread(Thorne et al., 2021). We propose that a detailed understanding of the innate immune mechanisms that protect from zoonosis, and pandemic spread after spillover, will be critical for management of future pandemic risk and propose that the well understood HIVs offer excellent tools for further relevant discovery in this field.

## SUPPLEMENTARY FIGURE LEGENDS

**Supplementary Figure 1.**
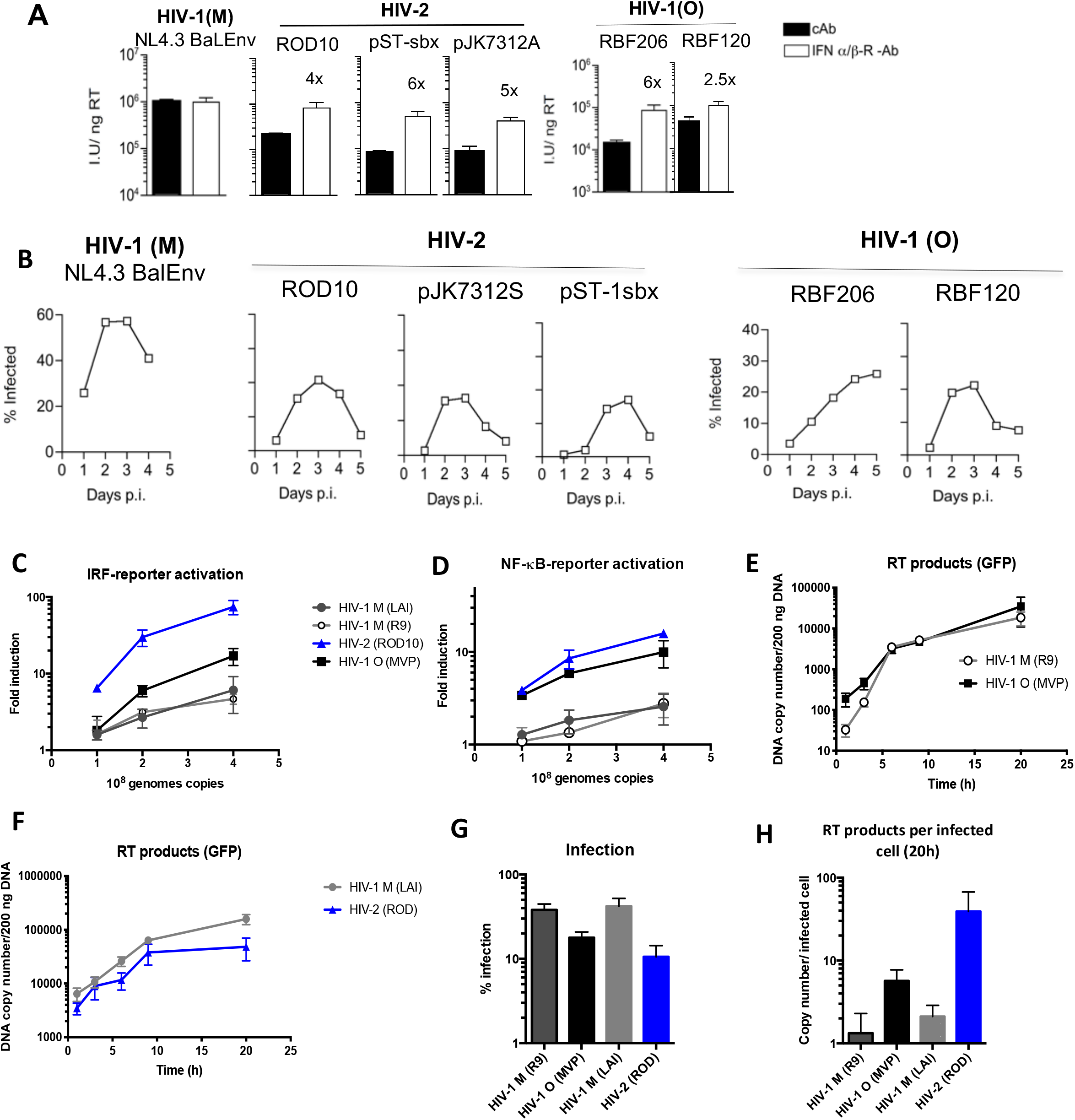
**(A)** Infection of MDM with HIV, measured at 48h by counting Gag positive cells, in the presence of anti-interferon α/β receptor (IFNα/β-R), or control, antibody (cAb). (**B)** Replication of HIV-1(M), HIV-2 or HIV-1(O) isolates in permissive GHOST cells measuring induced GFP expression by flow. **(C, D)** Activation of **(C)** IRF-luciferase reporter or **(D)** NF-kB secreted alkaline phosphatase reporter 48h after infection by equal genome copies of VSV-G-pseudotyped HIV-1(M), HIV-2 or HIV-1(O) -GFP. **(E, F)** Measurement of VSV-G pseudotyped HIV-1(M), HIV-1(O) and HIV-2 DNA synthesis (GFP primers) during a 20h time course in THP-1 cells. (**G)** Infection measured at 48 hours in wells parallel to (e) by flow. (**H)** Viral DNA (GFP) copy number at 20h post-infection per infected cell using data from (E-F). Mean +/− SD, N=3 donors (MDM) or independent experiments (THP-1).

**Supplementary Figure 2.**
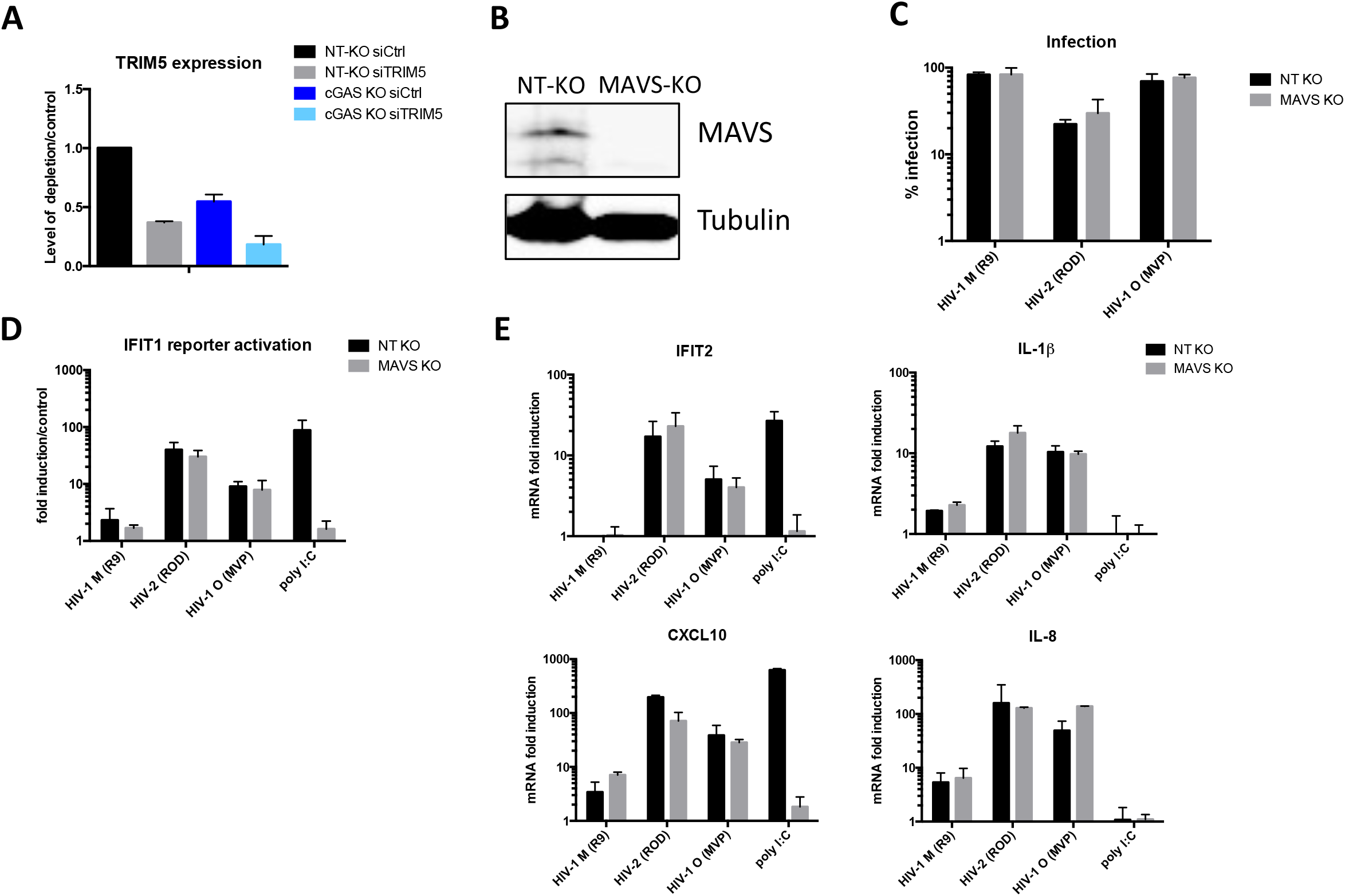
**(A)** GAPDH-normalised TRIM5 mRNA levels, measured after transfection of THP-1 treated with non-targeting CRISPR (NT-KO) or cGAS CRISPR (cGAS-KO) and transfected with TRIM5 targeting siRNA (siTRIM5) or non-targeting control siRNA (siCtrl). **(B)** Anti-MAVS or tubulin western blot of non-targeting KO cells (NT-KO) or MAVS KO cells. **(C)** % infection of HIV-1 (M), HIV-2 and HIV-1 (O) in NT-KO or MAVS KO cells. **(D)** IFIT1 reporter activation after viral infection normalised against mock infected. **(E)** GAPDH-normalised mRNA levels expressed as fold induction over mock-treated non-targeting KO control (NT-KO) or MAVS KO THP-1 cells 24h post-infection or after poly I:C transfection (500 ng/mL). Mean +/− SD, n= 3.

**Supplementary Figure 3.**
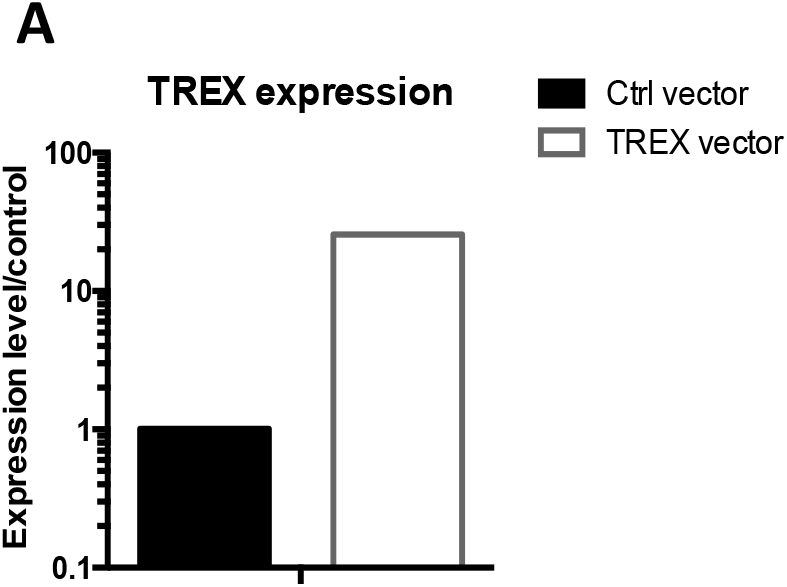
**(A)** A representative example of GAPDH-normalised TREX1 mRNA levels measured after transduction of THP-1 cells with empty (Ctrl) MLV vector or TREX1 expression vector.

**Supplementary Figure 4.**
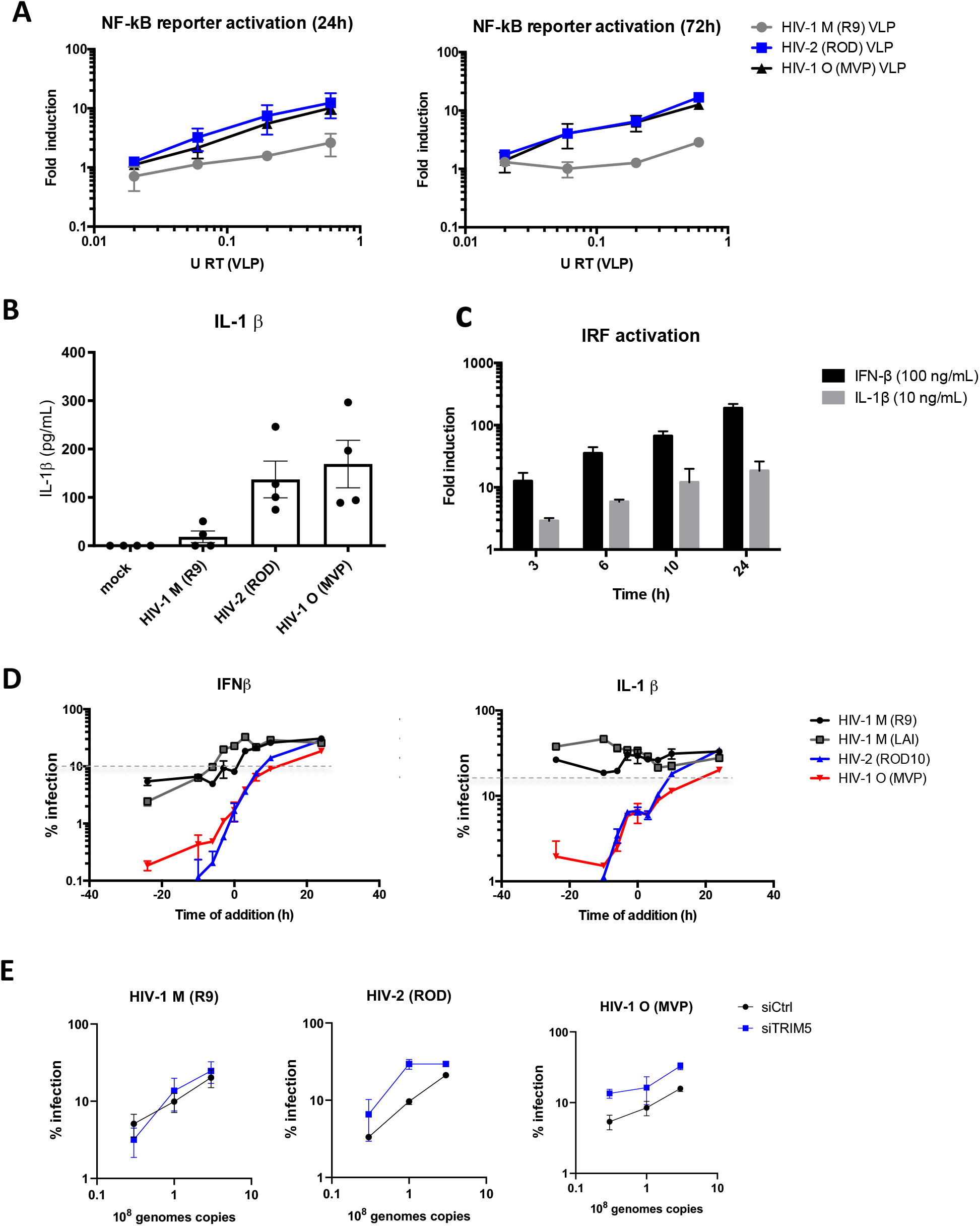
**(A)** NF-kB - secreted alkaline phosphatase reporter 24 or 72 hours after treatment with increasing doses of viral-like particles (VLP, made with packaging plasmid and VSV-G but no genome) (**B)** Quantification of IL-1β in supernatants from MDM mock infected or infected with HIV-1(M), HIV-2 or HIV-1(O). **(C)** IRF-reporter activation after interferon β (IFNβ) or IL-1β treatment of THP-1 cells. **(D)** % of infection in THP-1 cells after addition of IFNβ or IL-1β at different time points. Dotted line indicates the % of infection of untreated cells. **(E)** Infection levels in MDM depleted of TRIM5 with equal genome copies of VSV-G pseudotyped HIV-1(M), HIV-2 or HIV-1(O) –GFP measured 48h post-infection. Mean + SD, n= 3 independent experiments (ThP1) or 4 donors (MDM).

**Supplementary Figure 5.**
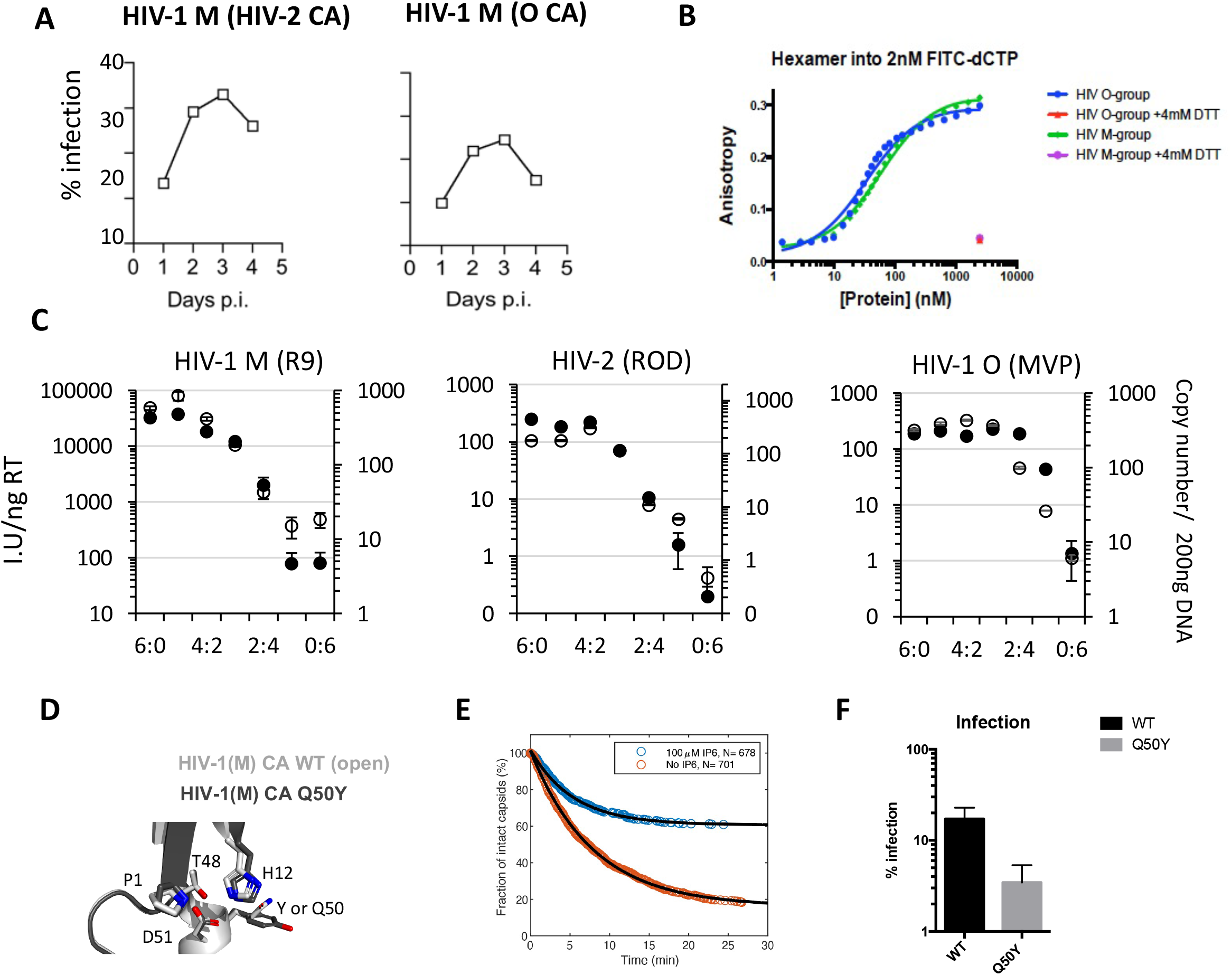
**(A)** Replication of HIV-1(M) NL4.3 (Bal Env) bearing HIV-2 ROD10 or HIV-1(O) MVP5180 Capsid in permissive GHOST cells (flow cytometry for induced GFP expression) representative of 3 independent experiments. **(B)** Binding of fluorescently labeled nucleotides to HIV-1(M) or HIV-1(O) recombinant CA hexamers in the presence or absence of DTT to reduce monomer cross links. **(C)** Titres of HIV-1(M), HIV-2 or HIV-1(O) –GFP made by mixing WT and CA R18G bearing packing constructs (left axis, white circles) and DNA synthesis measured at 6 hours post infection (right axis, black circles). **(D)** Amino acids in BHP hinge region influencing BHP position with overlay of HIV-1(O) (PDB ID:7T12) and open HIV-1(M) (PDB ID:5HGL) hexamers. **(E)** Capsid survival curves for CA mutant Q50Y generated from pooled data from two (no IP6) or three (100 μM IP6) independent experiments showing IP-mediated capsid stabilisation. **(F)** Single round infection of MDM with equal genome copies of VSV-G-pseudotyped HIV-1(M) (R9) CA Q50Y –GFP measured 48h post-infection by flow. Mean +/− SD, n= 3. (C, F)

**Supplementary Figure 6.**
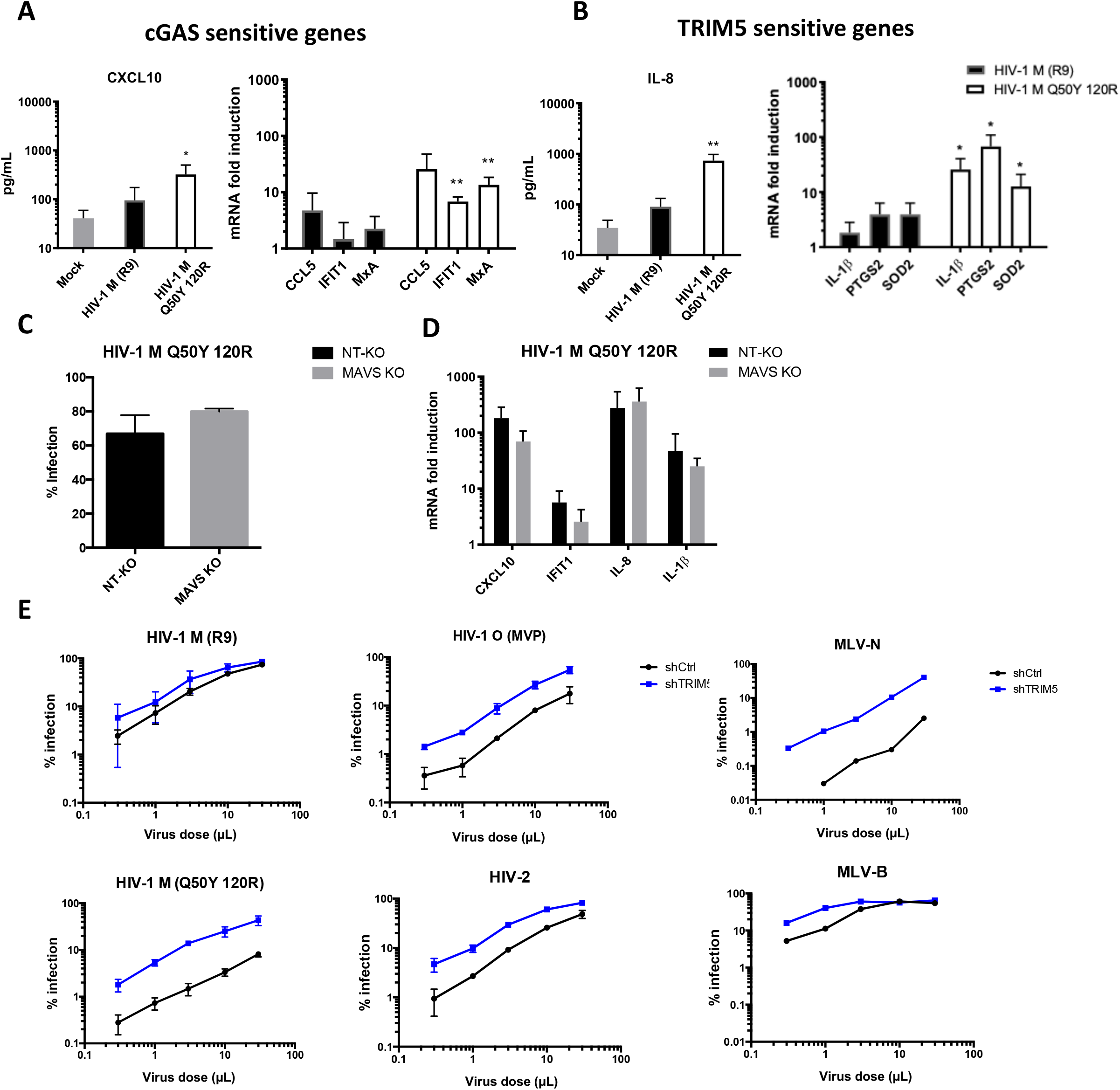
**(A, B)** Secreted CXCL10 (A) IL8 (B) (ELISA) 48hpi and GAPDH-normalised mRNA levels in MDM (fold induction over uninfected 24hpi. **(C)**% Infection of HIV-1(M) CA Q50Y 120R in non-targeting CRISPR (NT-KO) or MAVS CRISPR knock out (MAVS-KO) cells. **(D)** GAPDH-normalised mRNA levels, (fold induction over uninfected), in non-targeting CRISPR control (NT-KO) or MAVS KO THP-1 cells 24hpi. Mean +/− SD, n=3 independent experiments or donors. A, B unpaired t-test vs untreated MDM. \**p*<0.05, \*\**p*<0.01, **(E)** VSV-G pseudotype titration curves in U87 cells expressing non-targeting control shRNA (shCtrl) or TRIM5 targeting shRNA (shTRIM5). Mean +/− SD n= 3.

## METHODS

### Cells and reagents

HEK293T and U87 cells were maintained in DMEM (Gibco) supplemented with 10 % foetal bovine serum (FBS, Labtech) and 100 U/ml penicillin plus 100 μg/ml streptomycin (Pen/Strep; Gibco). THP-1-IFIT-1 cells that had been modified to express Gaussia luciferase under the control of the *IFIT-1* promoter were described previously (Mankan et al., 2014). THP-1 Dual Control and cGAS−/− cells were obtained from Invivogen. THP-1 IFIT1 cells were maintained in RPMI (Gibco) supplemented with 10 % FBS and Pen/Strep. THP-1 Dual cells were maintained in RPMI (Gibco) supplemented with 10 % FBS, Pen/Strep, 25 mM HEPES (sigma), 10 μg/ml of blasticidin (Invivogen) and 100 μg/ml of Zeocin™ (Invivogen). GHOST cells stably expressing CD4, CCR5, CXCR4, and the green fluorescent protein (GFP) reporter gene under the control of the HIV-2 long terminal repeat, were maintained in DMEM supplemented with 10% FBS, and antibiotics, G418 (500 μg/ml) (Thermo Fisher), hygromycin (100 μg/ml)(Invitrogen) and puromycin (1 μg/ml) (Millipore). Lipopolysaccharide, IFNβ, IL-1β and poly I:C were obtained from Peprotech. Herring-testis DNA was obtained from Sigma. For stimulation of cells by transfection, transfection mixes were prepared using lipofectamine 2000 in Optimem (Thermofisher Scientific) according to the manufacturer’s instructions (Invitrogen). HT DNA and poly I:C concentration used are stated on each figure.

### Isolation of primary monocyte-derived macrophages

Primary monocyte-derived macrophages (MDM) were prepared from fresh blood from healthy volunteers. The study was approved by the joint University College London/University College London Hospitals NHS Trust Human Research Ethics Committee and written informed consent was obtained from all participants.

Peripheral blood mononuclear cells (PBMCs) were isolated by density gradient centrifugation using Lymphoprep (Stemcell Technologies). PBMCs were washed three times with PBS and plated to select for adherent cells. Non-adherent cells were washed away after 1.5 h and the remaining cells incubated in RPMI (Gibco) supplemented with 10 % heat-inactivated pooled human serum (Sigma) and 100 ng/ml macrophage colony stimulating factor (R&D systems). For replication experiments with full-length viruses, the medium was then refreshed after 3 days (RPMI 1640 with 10% human serum), removing any remaining non-adherent cells. After 6 days, media was replenished with RPMI containing 5% type AB human serum (Sigma-Aldrich). For single-round experiments with VSV-G-pseudotyped viruses, on day 3 of differentiation cells were further washed and the medium changed to RPMI supplemented with 10 % heat-inactivated FBS. MDM were then infected 3-4 days later. Replicate experiments were performed with cells derived from different donors.

### Editing of cells by CRISPR/Cas 9

Lentiviral particles to generate CRISPR/Cas9-edited cell lines were produced by transfecting 10 cm dishes of HEK293T cells with 1.5 μg of pLentiCRISPRv2 encoding gene specific guide RNAs (Addgene plasmid #52961), 1 μg of p8.91 packaging plasmid(Zufferey et al., 1997), and 1 μg of vesicular stomatitis virus-G glycoprotein expressing plasmid pMDG (Genscript) using Fugene 6 transfection reagent (Promega) according to the manufacturer’s instructions. Virus supernatants were harvested at 48 and 72 h post-transfection, pooled and used to transduce THP-1 IFIT-1 cells by spinoculation (1000 *xg*, 1 h, room temperature). Transduced cells were selected using puromycin (1 μg/ml, Merck Millipore) and single clones isolated by limiting dilution in 96 well plates. Clones were screened for successful gene knock out by luciferase assay after targeted protein stimulation and immunoblotting. gRNA sequences: MAVS: CAGGGAACCGGGACACCCTC Non-targeting control: ACGGAGGCTAAGCGTCGCAA

### Virus plasmids

The NL4.3 (Ba-L Env), and YU2 molecular clones have been described(Rasaiyaah et al., 2013). O group molecular clones, RBF206 and BCF120, and HIV-2 molecular clones pJK7312S(Gao et al., 1992), pST(Kumar et al., 1990) have been described. Molecular clone ROD10 was obtained from National Institute of Biological Standards and Controls AIDS reagents programme(Clavel et al., 1986). The CA chimera molecular clone was generated by overlap PCR, replacing CA residues 1–204 of NL4.3 with the equivalent residues from MVP5180 or HIV-2 ROD10. VSV-G pseudotyped GFP-encoding vectors include HIV-1 M LAI ΔEnv.GFP (LAI strain(Peden et al., 1991)) with the Nef coding region replaced by GFP, HIV-1(M) R9 packaging vector (p8.91) and minimum genome expressing GFP (CSGW)(Bainbridge et al., 2001). HIV-2 ROD GFP has been described(Griffin et al., 2001). HIV-1(O) packaging plasmid to make HIV-1(O) GFP was generated by replacing Gag-Pro residues between *Not*1–*Bcl*1 in p8.91 with the equivalent residues from MVP5180(Ikeda et al., 2004). Q50Y, 120R mutations were generated by site directed mutagenesis of p8.91using *Pfu*Turbo (Agilent Technologies). For TREX1 overexpression we used MLV-based gammaretroviral expression vector EXN(Zhang et al., 2006), where TREX1 coding sequence was cloned from a plasmid kindly provided by Nan Yan between *BamHI* and *XhoI* sites. For TRIM5 depletion with shRNA, we expressed shRNA using SIREN-RetroQ (Clontech) gammaretroviral vector containing shRNA sequence targeting human TRIM5(Stremlau et al., 2004) or scramble Ctrl(Fletcher et al., 2015) as described.

### Production of virus in HEK293T cells

Replication competent HIV were produced by transfection of HEK293T cells in T150 flasks using Fugene 6 transfection reagent (Promega) according to the manufacturer’s instructions. Briefly, just-subconfluent T150 flasks were transfected with 8.75 μg of vector and 30 μl Fugene 6 in 500 μl Optimem (Thermofisher Scientific). Virus supernatants were collected 48 h, 72 h and 96 h post transfection. Virus suspensions were filtered, subject to ultracentrifugation through a 20% sucrose buffer and resuspended in RPMI 1640 with 5% HS for subsequent replication experiments in MDM. For VSVG-pseudotyped GFP expressing virus, each T150 flasks was transfected with 2.5 μg of vesicular stomatitis virus-G glycoprotein expressing plasmid pMDG (Genscript) and 6.25 μg of pLAIΔEnv GFP or 2.5 μg packaging plasmid (p8.91, MVP or HIV-2-pack) and 3.75 μg of GFP encoding genome plasmid (CSGW or HIV-2 GFP) using 30 μl Fugene 6 in 500 μl Optimem. In the case of viral-like particles (VLP) without genome, the cells in T150 flasks were transfected only with 2.5 μg of vesicular stomatitis virus-G glycoprotein expressing plasmid pMDG and 5 μg packaging plasmid (p8.91, MVP or HIV-2). Virus supernatants were harvested at 48 and 72 h post-transfection, pooled, DNase treated (2 h at 37°C, DNaseI, Sigma) and subjected to ultracentrifugation over a 20 % sucrose cushion. Viral particles were finally resuspended in RPMI supplemented with 10 % FBS.

Lentiviral particles to generate TREX expressing vector or TRIM5 shRNA vector were produced by transfecting 10 cm dishes of HEK293T cells with 1.5 μg of TREX EXN vector or TRIM5 targeting SIREN-RetroQ, 1 μg of packaginf plasmid CMVintron, and 1 μg of vesicular stomatitis virus-G glycoprotein expressing plasmid pMDG using Fugene 6 transfection reagent according to the manufacturer’s instructions. Virus supernatants were harvested at 48 and 72 h post-transfection, pooled and stored at −80°C.

### Virus quantification and RT products

Full-length HIV clones were quantified by reverse transcriptase (RT) enzyme-linked immunosorbent assay (ELISA) (Roche) according to manufacturer’s instructions. Reverse transcriptase activity of virus preparations was quantified by qPCR using a SYBR Green-based product-enhanced RT (SG-PERT) assay as described (Vermeire, Naessens et al., 2012). For viral genome copy measurements, RNA was extracted from 2 μl sucrose purified virus using the RNeasy mini kit (QIAgen). The RNA was then treated with TURBO DNase (Thermo Fisher Scientific) and subjected to reverse transcription using Superscript III reverse transcriptase and random hexamers according to the manufacturer’s protocol (Invitrogen). Genome copies were then measured by Taqman qPCR using primers against GFP(Towers et al., 1999) (see below).

For RT product measurements DNA was extracted from 5×10^5^ infected cells using the DNeasy Blood & Tissue kit (QIAgen) according to the manufacturer’s protocol. DNA concentration was quantified using a Nanodrop for normalisation. RT products were quantified by Taqman qPCR using TaqMan Gene Expression Master Mix (ThermoFisher) and primers and probe specific to GFP. A dilution series of plasmid encoding GFP was measured in parallel to generate a standard curve to calculate the number of GFP copies.

GFP primers:

*GFP* fwd: 5’- CAACAGCCACAACGTCTATATCAT -3’
*GFP* rev: 5’- ATGTTGTGGCGGATCTTGAAG -3’
*GFP* probe: 5’- FAM-CCGACAAGCAGAAGAACGGCATCAA-TAMRA -3’

### Infection assays

To measure viral replication, MDM were infected with 100pg RT of full-length viruses, measured by RT ELISA (Roche), per well (MOI 0.2) in 48-well plates and subsequently fixed and stained using CA-specific antibodies (EVA365 and EVA366 National Institute of Biological Standards AIDS Reagents Programme) and a secondary antibody linked to beta galactosidase, as described (Rasaiyaah et al., 2013). Anti-IFN-α/β receptor (PBL Interferon Source) or control IgG2A antibody (R&D systems) were added at 1 μg/ml for 2 h before infection and supplemented every 4 days. Single round-infection by VSVG-pseudotyped viruses was performed in 48-well plates using equal viral doses (1×10^9^ genome copies). Viral infection was measured 48 h later by enumeration of GFP positive cells by flow cytometry. For RNA extraction and subsequent qPCR analysis, cells were infected in 24-well plates.

GHOST cells were infected as described (Rasaiyaah et al., 2013) with full-length viruses, measured by RT ELISA, per well in 48-well plates. Cells were fixed at the indicated times post infection and GFP+ cells measured by flow cytometry.

Monocytic THP-1 cells were infected at a density of 2×10^5^ cells/ml in 48 well plates in the presence of polybrene (8 μg/ml, Sigma). Infection levels were assessed at 48 h post-infection through enumeration of GFP positive cells by flow cytometry. Input dose of virus was normalised either by RT activity (measured by SG-PERT) or genome copies (measured by qPCR) as indicated. THP-1 cells were treated with similar doses of viral-like particles (VLP) normalised by SG-PERT as indicated. After 24 h, cells were infected with equal amounts of genome copies (2×10^8^) and infection levels were measured 48 h post-infection through enumeration of GFP positive cells by flow cytometry. THP-1 cells stably expressing TREX were generated by transduction with the MLV-based gammaretroviral expression vector EXN. 4×10^4^ cells were spinoculated with 2 mL of Ctrl vector or TREX vector for 1 h at 1000g. These cells were maintained under selection with G418 (500 μg/ml).

100 ng/mL of IFNβ or 10ng/mL IL-1β were added at different time points to THP1-cells, which were then infected with a MOI of 0.3. Infection levels were measured after 48h by flow cytometry.

### Luciferase and secreted alkaline phosphatase reporter assays

Gaussia/Lucia luciferase activity was measured by transferring 10 μl supernatant to a white 96 well assay plate, injecting 50 μl per well of coelenterazine substrate (Nanolight Technologies, 2 μg/ml) and analysing luminescence on a FLUOstar OPTIMA luminometer (Promega). Data were normalised to a mock-treated control to generate a fold induction. Secreted alkaline phosphatase was measured using QUANTI-Blue™ according to manufacturer’s instructions, using 20 uL of cell supernatant.

### Quantitative RT-PCR

RNA was extracted from MDM or THP-1 cells using RNAeasy mini kit (Qiagen) according to the manufacturer’s instructions. 500 ng RNA was used to synthesise cDNA using Superscript III reverse transcriptase (Invitrogen), also according to the manufacturer’s protocol. cDNA was diluted 1:5 in water and 2 μl was used as a template for real-time PCR using SYBR® Green PCR master mix (Applied Biosystems) and QuantiStudio 5 Real-Time PCR machine (Applied Biosystems). Expression of each gene was normalised to an internal control (*GAPDH*) and these values were then normalised to mock-treated control cells to yield a fold induction. The following primers were used:

*GAPDH*: Fwd 5’-GGGAAACTGTGGCGTGAT-3’, Rev 5’-GGAGGAGTGGGTGTCGCTGTT-3’
*CXCL-10*: Fwd 5’-TGGCATTCAAGGAGTACCTC-3’, Rev 5’-TTGTAGCAATGATCTCAACACG-3’
*IFIT-2*: Fwd 5’-CAGCTGAGAATTGCACTGCAA-3’, Rev 5’-CGTAGGCTGCTCTCCAAGGA-3’
*MxA*: Fwd 5’-ATCCTGGGATTTTGGGGCTT-3’, Rev 5’-CCGCTTGTCGCTGGTGTCG-3’
*CCL5*: Fwd: 5’-CCCAGCAGTCGTCTTTGTCA-3’, Rev 5’-TCCCGAACCCATTTCTTCTCT-3’
*IFIT1*: Fwd: 5’-CCTCCTTGGGTTCGTCTACA-3’, Rev 5’-GGCTGATATCTGGGTGCCTA-3’
*IL-8*: Fwd: 5’-ATGACTTCCAAGCTGGCCGTGGCT-3’, Rev 5’-TCTCAGCCCTCTTCAAAAACTTCTC-3’*PTGS2*: Fwd: 5’- CTGGCGCTCAGCCATACAG-3’, Rev 5’-CGCACTTATACTGGTCAAATCCC-3’
*IL-1β*: Fwd: 5’-ATGATGGCTTATTACAGTGGCAA-3’, Rev 5’-GTCGGAGATTCGTAGCTGGA-3’
*SOD2*: Fwd: 5’-GGAAGCCATCAAACGTGACTT-3’, Rev 5’-CCCGTTCCTTATTGAAACCAAGC-3’
*cGAS*: Fwd 5’*-*GGGAGCCCTGCTGTAACACTTCTTAT-3’ Rev, 5’-TTTGCATGCTTGGGTACAAGGT-3’
*TREX*: Fwd 5’*-*CGCATGGGCGTCAATGTTTT-3’ Rev, 5’-GCAGTGATGCTATCCACACAGAA-3’

TRIM5 expression levels were measured using TaqMan Gene Expression Assay according to manufacturer’s instructions detecting TRIM5 (FAM dye-labelled, TaqMan probe ID no. Hs01552559_m1), or the housekeeping gene OAZ1 (FAM dye-labelled, primer-limited, TaqMan probe ID no. Hs00427923_m1).

### ELISA

Cell supernatants were harvested for ELISA at 48 h post-infection/stimulation and stored at − 20°C. CXCL-10 and IL-8 protein was measured using Duoset ELISA reagents (R&D Biosystems) according to the manufacturer’s instructions.

### cGAS and TRIM5 depletion by RNAi

1×10^5^ MDM differentiated in M-CSF for 4 days were transfected with 25 pmol of siRNA SMART pool against cGAS (L-015607-02-0005), TRIM5 (L-007100-00-0005) or non-targeting control (D-001810-10-05) (Dharmacon) using Lipofectamine RNAiMAX Transfection Reagent (Invitrogen). Transfection medium was replaced after 18h with RPMI 1640 medium supplemented with 10% FCS and cells cultured for additional 3 days before infection.

5×10^5^/mL THP-1 Dual cells were transfected with 35 pmol of siRNA SMART pool against cGAS, TRIM5 or non-targeting control (Dharmacon) using Lipofectamine RNAiMAX Transfection Reagent (Invitrogen). Transfection medium was replaced after 18h with RPMI 1640 medium supplemented with 10% FCS and cells were plated in 48-well plates and infected as indicated. To deplete TRIM5 in U87 cells, pSIREN-RetroQ expressing shRNA TRIM5 was transduced at an MOI ~ 1 and shRNA-expressing cells selected with 10 μg/mL puromycin.

TRIM5 and cGAS expression were quantified by qPCR normalised to OAZ1 and GAPDH, respectively, by the ΔΔCt method.

### Immunoblotting

For immunoblotting of THP-1 IFIT1 NT-Ctrl or KO cells, 2×10^6^ cells were lysed in lysis buffer containing 50 mM Tris pH 8, 150 mM NaCl, 1 mM EDTA, 10% (v/v) glycerol, 1 % (v/v) Triton X100, 0.05 % (v/v) NP40 supplemented with protease inhibitors (Roche), clarified by centrifugation at 14,000 x *g* for 10 min, and the supernatants were boiled for 5 min in 6X Laemmli buffer (50 mM Tris-HCl (pH 6.8), 2 % (w/v) SDS, 10% (v/v) glycerol, 0.1% (w/v) bromophenol blue, 100 mM β-mercaptoethanol) before separating on 12 % polyacrylamide gel. Proteins were transferred to a Hybond ECL membrane (Amersham biosciences) using a semi-dry transfer system (Biorad). Primary antibodies used: Mouse-anti-MAVS (Cell Signaling Technology), Mouse-anti-tubulin (Abcam). Primary antibodies were detected with goat-anti-mouse IRdye 800CW infrared dye secondary antibodies and membranes imaged using an Odyssey Infrared Imager (LI-COR Biosciences).

### Single molecule analysis

Single molecule analyses were carried out as described (Marquez et al., 2018).

### Phylogenetics

A dataset of representative HIV and SIV sequences from the CA region of *gag* were downloaded from the Los Alamos HIV-1 sequence database and aligned manually. The phylogeny was estimated from the nucleotide sequences using RAxML 8 (Stamatakis, 2014) with substitution model GTR + Gamma and rooted consistent with phylogenies that include non-primate lentivirus outgroup taxa (Sharp and Hahn, 2011). ChromaClade was used to annotate taxon labels with residues found at capsid protein sites (Monit et al., 2019). Note that chromaclade does not use statistical tests to assess viral evolution. Rather it provides a simple way to visualise lineage specific amino acid variation in a very qualitative and intuitive way. In this study our focus on positions CA 50 and 120 was also influenced by the hexamer structures which revealed that these positions have a role in the structural differences observed between viral capsid hexamers.

### Protein production and purification

#### HIV-1 (M) CA R120

Protein was expressed and purified as previously described for HIV-1 (M) CA WT (Lanman et al., 2002). In brief, the plasmid encoding HIV-1 (M) CA R120 were transformed into *E. coli* C41 OverExpress™ C41(DE3) (Lucigen) following the manufacturer’s instructions. Transformed cells were plated onto LB agar plates supplemented with 100 μg/ml Ampicillin. A single colony were used to inoculate a 5 mL overnight starter culture consisting of 2YT media supplemented with 100 μg mL−1 Ampicillin. The culture were grown at 37°C in an orbital shaker at 250 rpm. The next day the starter culture were used to inoculate 1 L 2YT media supplemented with 100 μg/mL Ampicillin. The culture were grown at 37°C in an orbital shaker at 250 rpm until OD600 nm reached 0.5, followed by the addition of 0.4 mM IPTG to induce overexpression. Protein expression was carried out overnight at 14°C. After expression the cells were harvested by centrifugation at 6000×g for 20 min followed by resuspension in Lysis Buffer (50 mM Tris-HCl, 40 mM NaCl, 20 mM β-mercaptoethanol [pH 4.5]), lysed using a cell disruptor, followed by removal of the insoluble fraction by centrifugation at 30000 × g for 20 min. The capsid protein was precipitated by adding 20 % (w/v) ammonium sulphate and pelleted by centrifugation at 30000 × g for 20 min. The obtained pellets were resuspended in Refolding Buffer (100 mM citric acid, 20 mM [pH 4.5]). This was followed by extensive dialysis against the same buffer. Lastly, the protein was dialysis against 25 mM Tris-HCl [pH 8]. The capsid was further purified with anion exchange chromatography (AEC) using a 5 mL Hi-TRAP Q column (Cytiva). The AEC purification was performed using buffer A (25 mM Tris-HCl [pH 8]) and Buffer B (25 mM Tris-HCl, 1M NaCl [pH 8]). Lastly SEC was performed using a Superdex 16/600 75 pg column and a buffer containing 25 mM Tris-HCl and 40 mM NaCl [pH 8]. All steps were performed on an ÄKTA pure system (Cytiva). The protein was further concentrated to 3.0 mg/mL in the SEC buffer for crystallisation.

#### HIV-1(O), HIV-1(M) CA Q50Y, SIVmac, SIVcpz

Hexameric CA proteins, stabilized by engineered inter-subunit disulfide bonds, were produced by assembly of recombinant CA containing four amino acid substitutions, as previously reported (Pornillos et al., 2010). Briefly, these CA mutations are: HIV-1 (O-group) (A14C, E45C, W185A, M186A); HIV-1 (M-group) (A14C, E45C, Q50Y, W184A, M185A); SIVmac (P13C, E44C, W182A, M183A); and SIVcpz (P14C, E45C, W184A, M185A). All hexameric assemblies were expressed and purified as previously described (Jacques et al., 2016). Briefly, expression performed in *E. coli* (C41) by mid-log induction with 1 mM IPTG. Induced cells were incubated overnight at 14 °C and harvested by centrifugation. Cells were lysed in 50 mM TRIS (pH 8.0), 150 mM NaCl, 20 mM β-mercaptoenthanol by sonication. Clarified lysates were treated with 20 % (w/v) ammonium sulfate, and the precipitate resuspended in 100 mM citric acid (pH 4.5), 20 mM β-mercaptoenthanol and dialysed against the same to remove the ammonium sulfate. Redissolved protein was subjected to assembly by three dialysis steps: (1) 1 M NaCl, 50 mM TRIS (pH 8.0), 20 mM β-mercaptoenthanol; (2) 1 M NaCl, 50 mM TRIS (pH 8.0); and (3) 20 mM TRIS (pH 8.0, 40 mM NaCl. Purified hexamers were isolated by size-exclusion chromatography using a 16/600 Superdex 200 Prep Grade column pumped on an ÄKTA Pure with 20 mM TRIS, 40 mM NaCl.

### Crystallisation, structure solution and analysis

#### HIV-1 (M) CA R120

Crystals were grown using the hanging-drop vapour diffusion technique at 20°C by mixing 1 μL protein with 1 or 2 uL precipitant, containing 9.5-11% (w/v) PEG3350, 250-350 mM NaI, and 100 mM Sodium Cacodylate [pH 6.5] as previously described (Gres et al., 2015). Crystals of hexagonal shape appeared after 3-5 days reaching a maximum size of 0.1 – 0.2 mm after 2 weeks. Before data collection, the harvested crystals were immersed in a solution containing the precipitant mixture and 20 % (v/v) glycerol and cryo-cooled in liquid nitrogen. Diffraction data were collected from a single crystal at the PETRA III P13 beamline (EMBL-Hamburg/DESY P13, Germany). The data set was indexed, processed, and scaled using the XDS package (Kabsch, 2010). The HIV-1 (M) CA R120 crystal belonged to the P 6 space group with a solvent content of 48.5% corresponding to one molecule per asymmetric unit. The structure was determined by molecular replacement using Phenix Phaser (Adams et al., 2010) and a previously determined HIV-1 (M) CA structure (PDB ID:4XFX) as search model. Model building was performed using COOT (Emsley et al., 2010). Refinement was performed using REFMAC 5.8. Overview of refinement procedures within REFMAC5: utilizing data from different sources (Nicholls et al., 2018) using a TLS/maximum likelihood protocol. The model converged to a final Rwork/Rfree of 0.242/0.277 at a resolution of 2.30 Å. The HIV-1 (M) CA R120 model covers the HIV-1 (M) CA amino-acid sequence 1-222 and contains in addition 2 Iodine, 4 Chlorine ions and 14 water molecules. Final figures were rendered in The PyMOL Molecular Graphics System, Version 2.5.0.a0 Schrodinger, LLC.

#### HIV-1(O), HIV-1(M) CA Q50Y, SIVmac, SIVcpz

HIV-1 (O-group) hexamer crystals were grown using hanging drop vapor-diffusion using 2 μl protein (40 mg/ml) + 2 μl crystallant (10% (w/v) PEG 6000, 100 mM HEPES (pH 7.0), 100 mM Glycine) suspended over 500 μL undiluted crystallant. Crystals were highly-susceptible to damage from standard cryoprotectants, and were cryoprotected with the gradual addition of glucose (solid) to a final concentration of 40% (w/v). HIV-1 (M-group, Q50Y) hexamer crystals were grown using hanging drop vapor-diffusion using 2 μl protein (13 mg/ml) + 2 μl crystallant (19% (v/v) PEG 550MME, 100 mM TRIS (pH 8.0), 150 mM KSCN, 10 mM ATP, 3% (v/v) 3,5-hexanediol) suspended over 500 μL undiluted crystallant. Crystals were cryoprotected in 20% (v/v) MPD. SIVmac hexamer crystals were grown using sitting drop vapor-diffusion using 200 nL protein (12 mg/ml) + 200 nL crystallant (10% (w/v) PEG 6000, 5% (w/v) MPD, 100 mM HEPES (pH 7.5)) suspended over 80 μL crystallant. Crystals were cryoprotected in 20% (v/v) MPD. SIVcpz hexamer crystals were grown using sitting drop vapor-diffusion using 200 nL protein (12 mg/ml) + 200 nL crystallant (4.5% PEG 550MME, 0.15M KSCN, 0.1M Tris (pH 9.0), 4% 2,5-hexanediol) suspended over 80 μL crystallant. Crystals were cryoprotected in 20% (v/v) MPD. Diffraction data were collected at 100 K on Diamond Light Source beamlines I02 (HIV-1(O-group) and HIV-1(M-group, Q50Y)), and I04-1 (SIVmac, SIVcpz). Data were reduced using IMOSFLM (Leslie and Powell, 2007) or XDS (Kabsch, 2010) and scaled and merged using AIMLESS (Evans and Murshudov, 2013). Structures were solved by molecular replacement using PHASER (McCoy, 2007) and search model based on the original crosslinked HIV-1 (M-group) hexamer, PDB:3H47 (Pornillos et al., 2010). Structures were refined using REFMAC5 (Murshudov et al., 1997) or phenix.refine (Afonine et al., 2012). Between rounds of refinement, models were manually checked and corrected against the corresponding electron-density maps in COOT (Emsley and Cowtan, 2004). The quality of the model was regularly checked for steric clashes, incorrect stereochemistry and rotamer outliers using MOLPROBITY (Chen et al., 2010).

### Statistical analysis

We have included number of replicates (equal to number of different donors), statistical tests, and significance criteria in figure legends and in the main text of the manuscript.

Statistical analysis was performed in GraphPad Prism. Following P-values were considered as significant: \*\*\**p*-value<*0.001, **p-*value ≤ 0.01, *\*p-*value ≤ 0.05

## Acknowledgements

SP and RJM were funded by UCL MRC doctoral training programme PhD studentships. LCJ is funded by the MRC (UK; U105181010), a Wellcome Trust Investigator Award (200594/Z/16/Z). FBR and BHH are funded by NIH RO1 AI120810. DAJ is funded by an Australian Research Council Discovery Project (DP180101384). DAJ and TB are funded by a National Health and Medical Research Council Project Grant (GNT1158338). GJT was funded by a Wellcome Trust Senior Biomedical Research Fellowship (108183) followed by a Wellcome Investigator Award (220863), the European Research Council under the European Union’s Seventh Framework Programme (FP7/2007-2013)/ERC (grant HIVInnate 339223) and the National Institute for Health Research University College London Hospitals Biomedical Research Centre. GJT, LCJ TB and DAJ are funded by a Wellcome Collaborative award (214344). We acknowledge Diamond Light Source for time on Beamlines I02 and I04-1 under Proposals mx8547-149 and mx11235-7. We acknowledge use of facilities in the Structural Biology Facility within the Mark Wainwright Analytical Centre– UNSW, funded in part by the Australian Research Council Linkage Infrastructure, Equipment and Facilities Grant: ARC LIEF 190100165. We also acknowledge access to the EMBL beamlines at PETRA III (DESY) supported by iNEXT-Discovery, project number 12462, funded by the European Commission Horizon 2020 program. Synchrotron data was collected at beamline P13 operated by EMBL Hamburg at the PETRA III storage ring (DESY, Hamburg, Germany) with thanks to Gleb Bourenkov for beamline assistance. We thank Nan Yan University of Texas at Austin for the TREX1 plasmid.

